# High-resolution modeling of thermal thresholds and multiple environmental influences on coral bleaching for regional and local reef managements

**DOI:** 10.1101/211854

**Authors:** Naoki H. Kumagai, Hiroya Yamano, Committee Sango-Map-Project

## Abstract

Corals are one of the communities most threatened by global and local stressors. Excessive summer sea temperatures can cause coral bleaching, resulting in decreases in living coral coverage. Coral bleaching may begin with rising sea temperatures, although the widely used threshold of 1 °C over the local climatological maximum sea temperature has been reconsidered. In this study, we refine thermal indices predicting coral bleaching at high resolution (1 km) by statistically optimizing the thermal threshold and multiple environmental influences on bleaching, such as ultraviolet (UV) radiation, water turbidity, and cooling effects on corals. We use a dataset of coral bleaching events observed during 2004–2016 in Japan derived from the Web-based monitoring system, the Sango (Coral) Map Project, aiming at regional to local conservation of Japanese corals. We show how the ability to predict coral bleaching is improved by the choice of thermal index, statistical optimization of thermal thresholds, usage of multiple environmental influences, and modeling methods (generalized linear model and random forest). After optimization, the differences among the thermal indices in the ability to predict coral bleaching were slight. Among environmental influences, cooling effects, UV radiation, and water turbidity, in addition to a thermal index, well explain the occurrence of coral bleaching. Prediction based on the best explanatory model reveals that recent Japanese coral reefs are experiencing bleaching in many areas, although we show a practical way to reduce bleaching frequency significantly by screening UV radiation. Thus, our high-resolution models may provide a quantitative basis for the management of local reefs under current global and local stressors. The results of this study may be useful to other researchers for selecting a predictive method according to their needs or skills.

## Introduction

Biological communities seem to shift toward alternative stable states in response to changing climate. Corals are likely to be one of the most susceptible communities to global warming and local environmental stressors (Hoegh-Guldberg, 1999; West & Salm, 2003).

Extreme rising sea temperature than in usual summer can cause a “bleaching” of reef-building corals. The excessive thermal stress leads to expulsion, digestion, or reduced pigmentation of symbiotic dinoflagellate algae in host coral cells, and the white skeleton of the corals becomes visible as if it were bleached (Hoegh-Guldberg, 1999; Brown et al., 2002). Prolonged warming trends in sea temperature will lead to increasingly frequent bleaching and mortality of corals in the future (Hoegh-Guldberg, 1999; Donner et al., 2005; Donner, 2009; McClanahan et al., 2015).

As an index representing cumulative thermal stresses, Degree heating weeks (DHW) by the National Oceanic and Atmospheric Administration Coral Reef Watch (NOAA CRW), has been widely employed as a predictive model of coral bleaching. DHW is based on sea-surface temperature (SST) derived from satellite images (Liu et al., 2003, 2014, 2017).

The index is the sum of excessive thermal stresses of 1 °C above historical summer monthly SST during 12 weeks (Fig. S1A). A DHW over 4 °C-weeks indicates coral bleaching (bleaching alert threshold; Fig. S1B). Despite its wide use, the predictive performance of DHW may be not sufficient for purposes of local reef management: DHW detects only 40% of global coral bleaching events (Donner, 2011). Its low performance may be due to its fixed thermal threshold of 1 °C above baseline SST. Indeed, many previous studies suggest that even thermal stress not exceeding 1 °C can induce coral bleaching (Brown, 1997; McWilliams et al., 2005; Kleypas et al., 2008) or historical temperature variability can affect bleaching (Brown et al., 2002; West & Salm, 2003). Therefore, some studies use modified indices such as the sum of thermal stresses over 0 °C above baseline SST, instead of 1 °C excess (e.g., Yee et al., 2008; Kayanne, 2017). Further, Donner (2011) proposed two modified DHWs: an index using historical SST variability as the bleaching alert threshold, and an index using the mean of the warmest monthly SST of each year instead of summer monthly SST as the baseline SST.

In addition to global measures to reduce climate warming, local measures to control environmental influences on coral resilience are needed for reef management (West & Salm, 2003). Prediction of where and when corals will bleach can be valuable information to support decision-making on managing local coral reefs. To evaluate global and local stressors on corals, a high-performance predictive model incorporating global and local stressors at high spatial resolution is required. Even global stressors such as thermal stresses can vary at the local scale (Strong et al., 2002; Liu et al., 2014). Further, there are potentially interacting environmental stressors, such as ultraviolet (UV) radiation (Hoegh-Guldberg, 1999; West & Salm, 2003; Maina et al., 2008; Yee et al., 2008) and variables that can reduce coral bleaching, including water turbidity, topography of the sea floor, and exposure to winds and currents (Hoegh-Guldberg, 1999; Nakamura & van Woesik, 2001; West & Salm, 2003; Oliver et al., 2009; Oxenford & Vallés, 2016).

Modeling coral bleaching at the local scale also requires high-resolution observational records at focal localities, because a low predictive model can be due to overlooking many bleaching events (Oliver et al., 2009). Observations in local areas of research interest have been well covered by the ReefBase (Tupper et al., 2011) and the Bleaching Database V1.0 (Donner et al., 2017). However, records from some areas are still limited in spite of much research interest. One possible reason for this is the language gap. A considerable amount of data in the databases may have been provided by nonprofessional (citizen) specialists who are not English speakers. Japan is an area from which bleaching records are hardly (*N ≤* 64) found in the global databases, despite much research interest.

Therefore, to collect and gather observational records of corals throughout Japan, diverse Japanese stakeholders associated with coral reefs, including professional scientists, constructed a Web-based monitoring system for Japanese coral reefs, the Sango Map Project, in 2008 (*sango* means coral in Japanese), the second International Year of the Reef (IYOR) (Namizaki et al., 2013). Collecting observational records via a Japanese Web-based database is quite effective in Japan, because internet service is available to the vast majority of people in Japan, who are able to use common Japanese much better than English, and most Japanese coral reefs develop around populated islands where diving services are available. The outcome of this project has been commented on in the IYOR international “Year in Review” report (Staub & Chhay, 2009).

In this study, we aimed to improve the ability to predict coral bleaching at high spatial resolution (1 km) sufficiently to contribute to regional and local reef management, using a dataset derived from the Sango Map Project. To this end, we compared the predictive performance of various thermal indices and their modifications, including models with multiple explanatory variables. As a part of the comparison, we developed another modification of DHW to represent less than 1 °C excess over baseline SST, using historical SST variability instead of the 1 °C threshold (hereafter “filtering threshold”) (Fig. S1C). Note that historical SST variability is globally less than 1 °C (Donner, 2011). Further, we optimized the filtering threshold by estimating statistically for each type of DHW and degree heating month (DHM) (Tables 1 and 2). Next, we optimized the combination of multiple explanatory variables together with optimizing the filtering threshold to maximize predictive performance. Finally, we produced maps of predicted coral bleaching in the main study areas and predictions under reduced local environmental stress as a guide to local reef management.

**Table 1.**
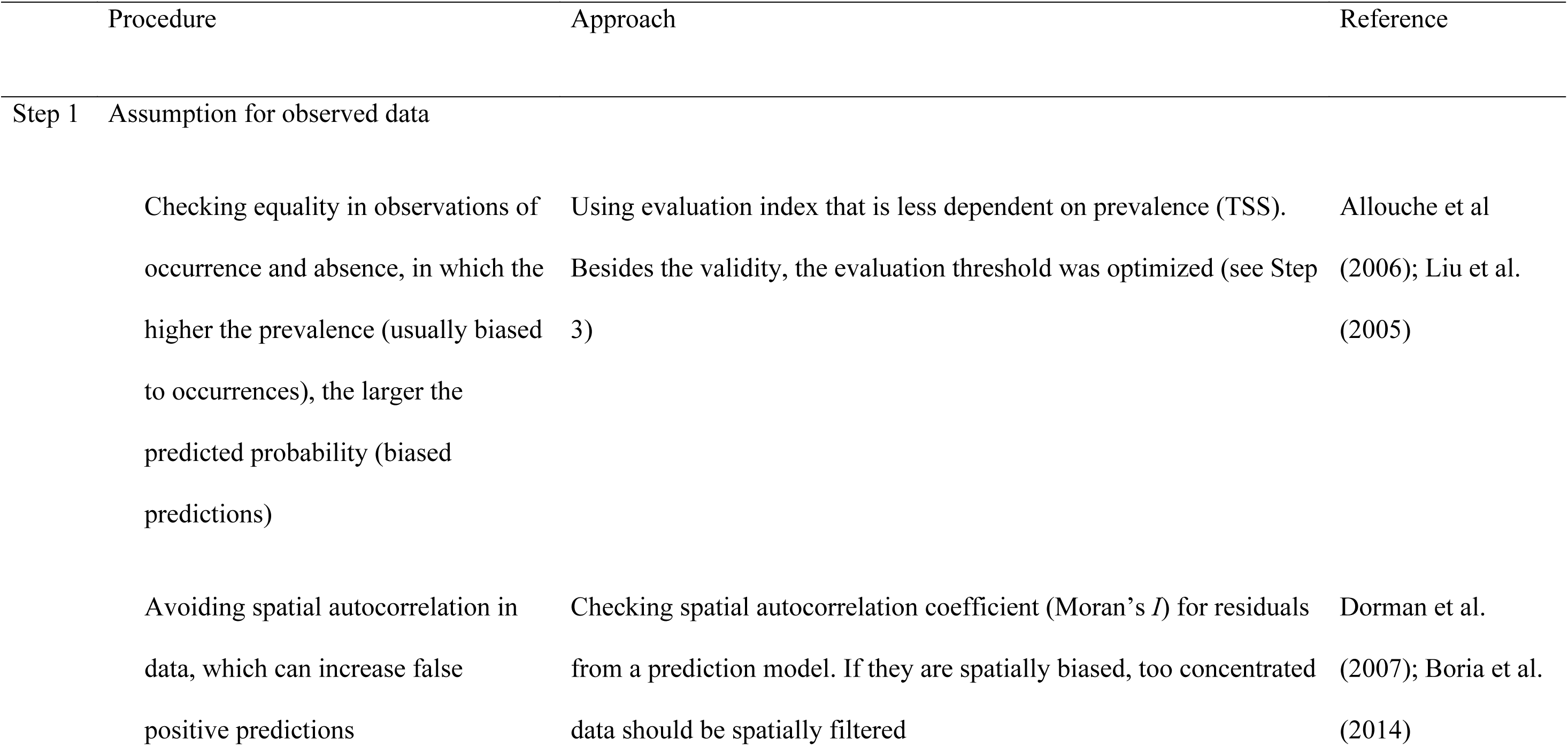
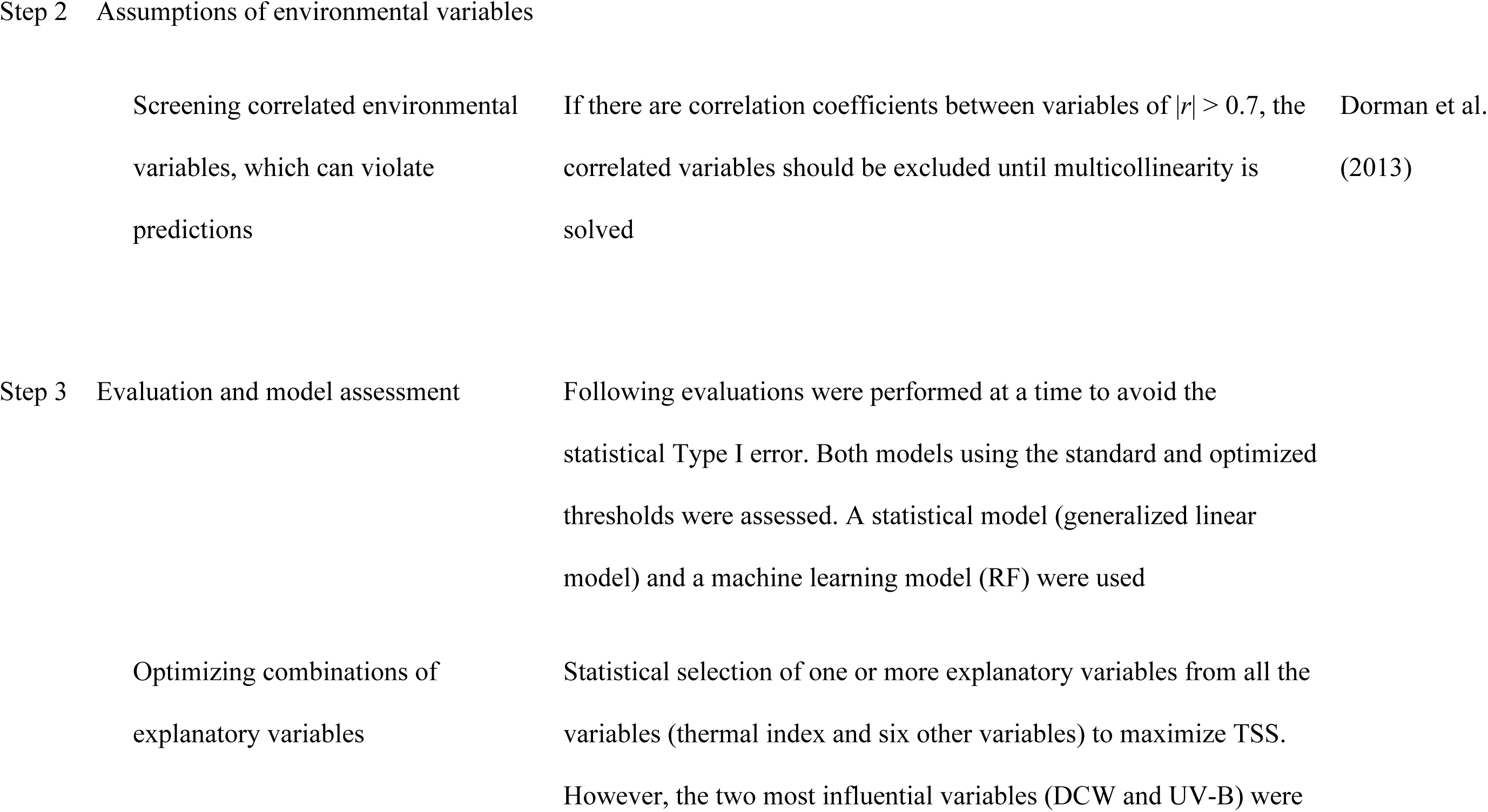
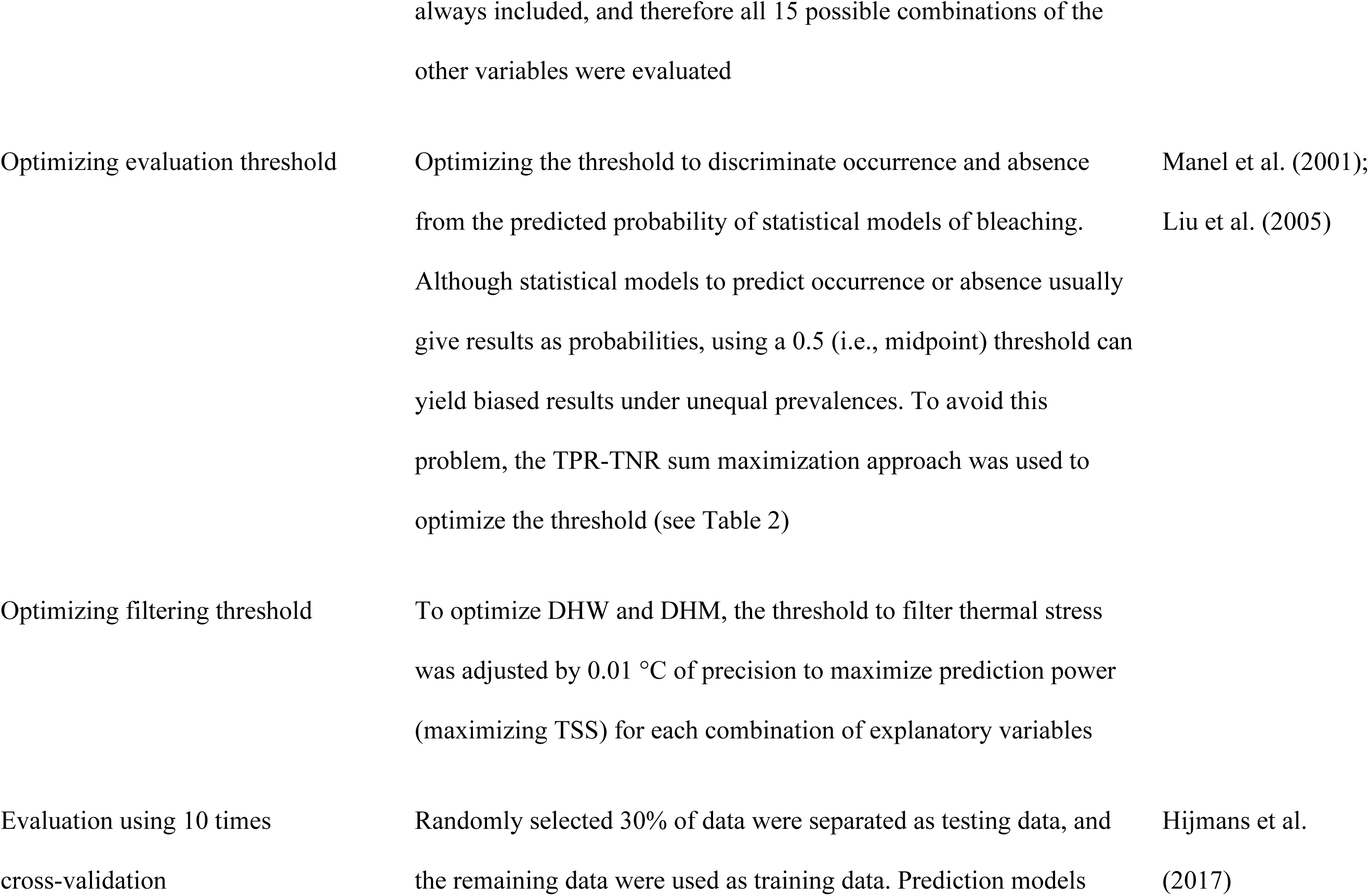
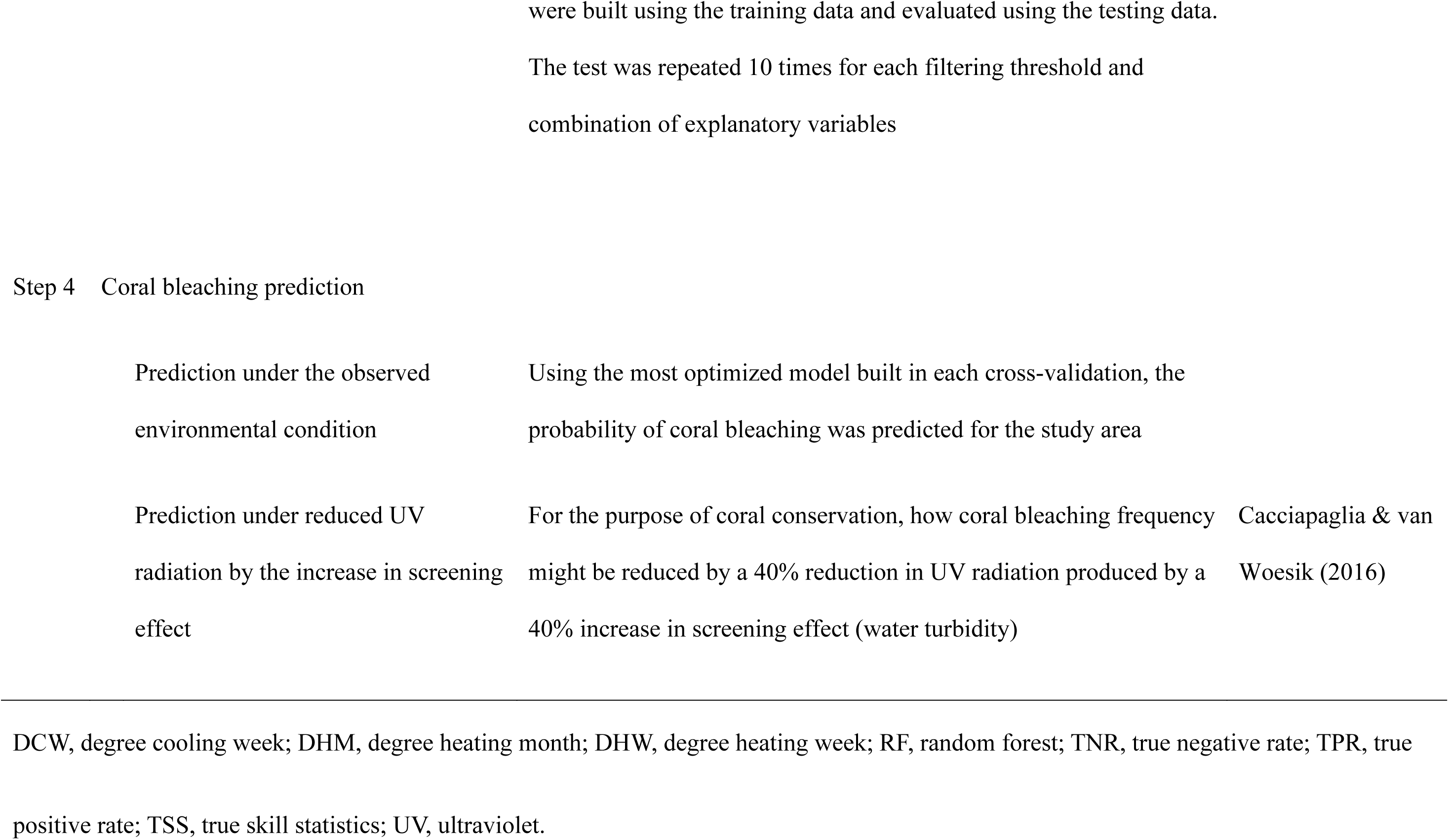
Flowchart summarizing the three steps in our analysis. Steps 1 and 2 confirmed the assumptions for explanatory variables and data, respectively. Then, Step 3 evaluated the prediction models. Finally, Step 4 performed predictions of coral bleaching.

**Table 2.**
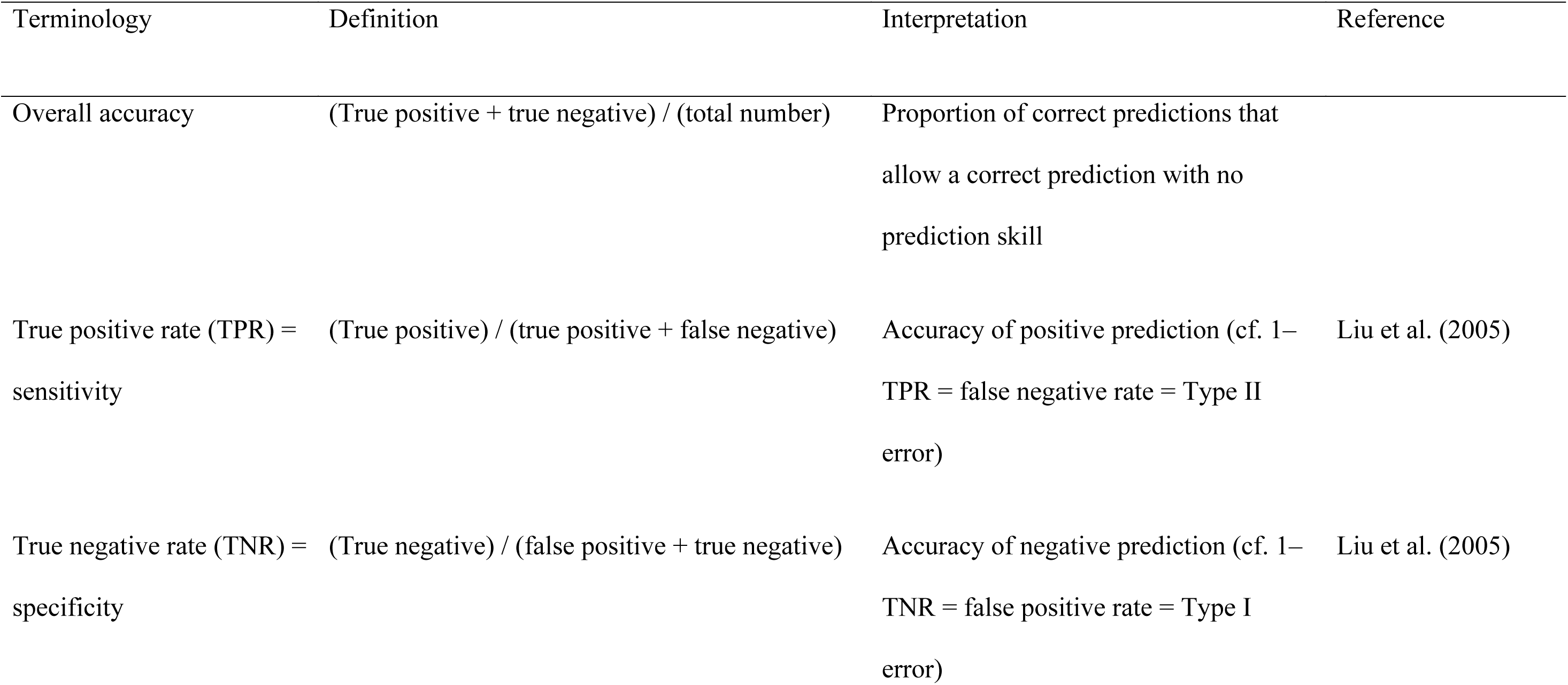
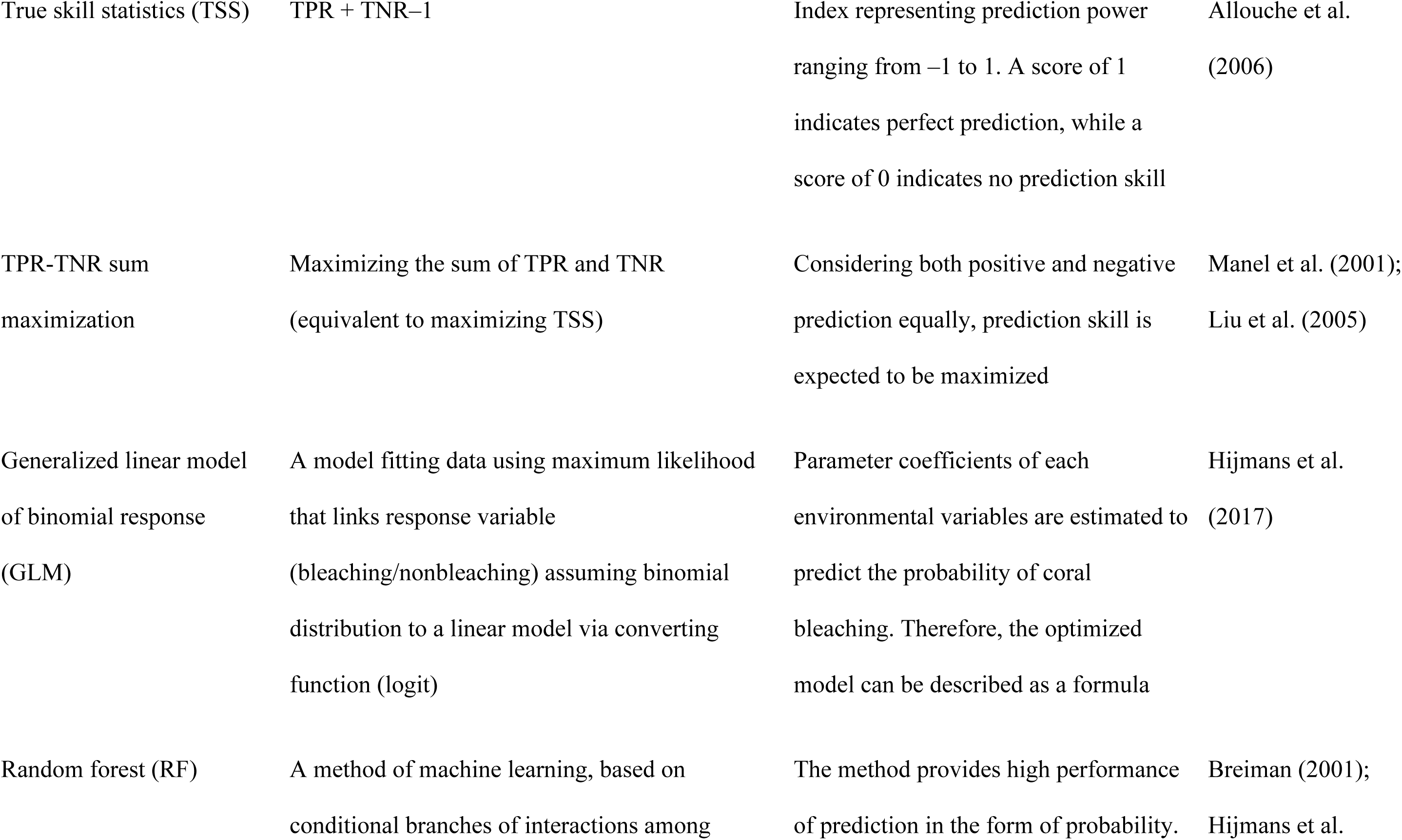
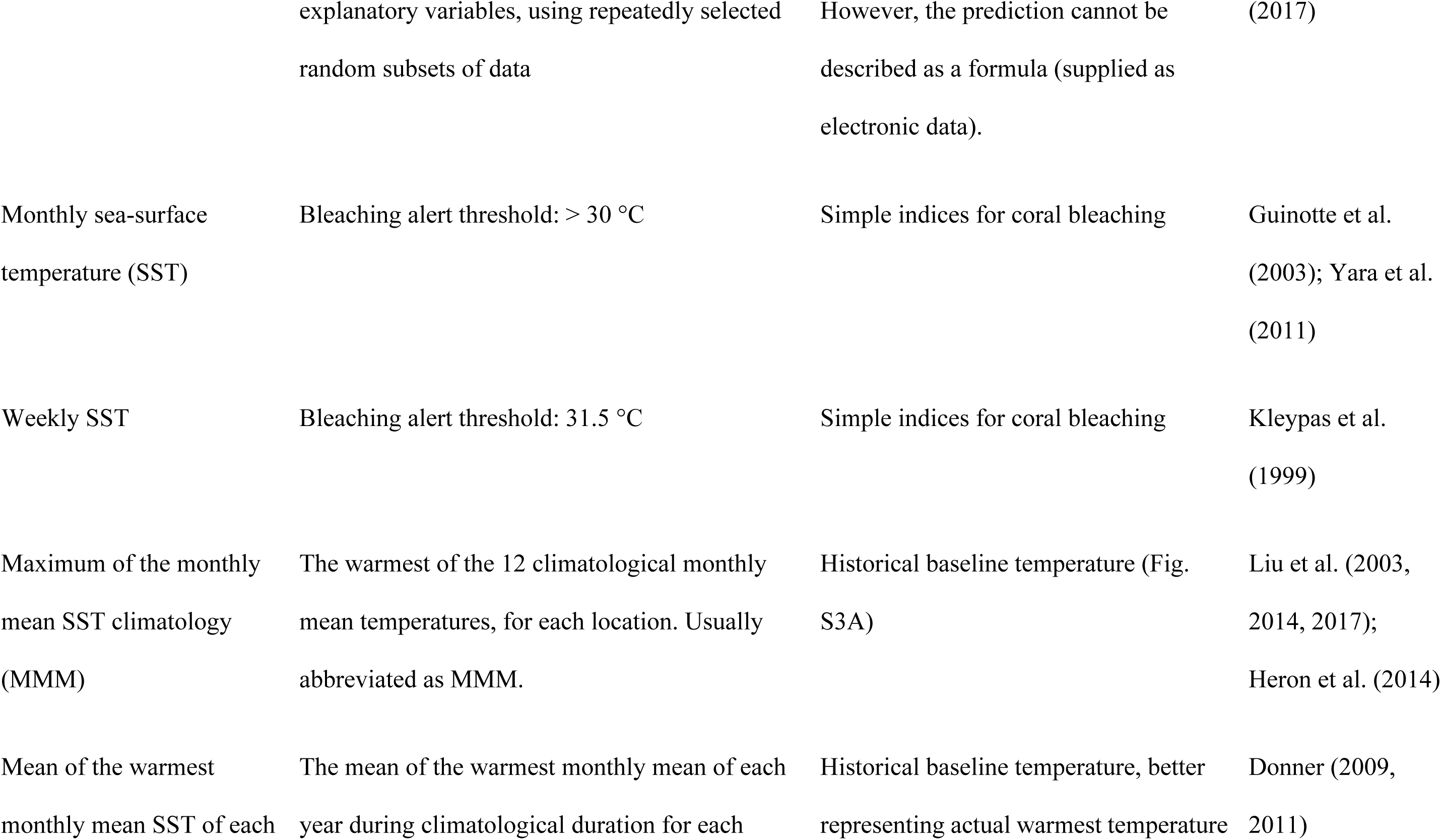
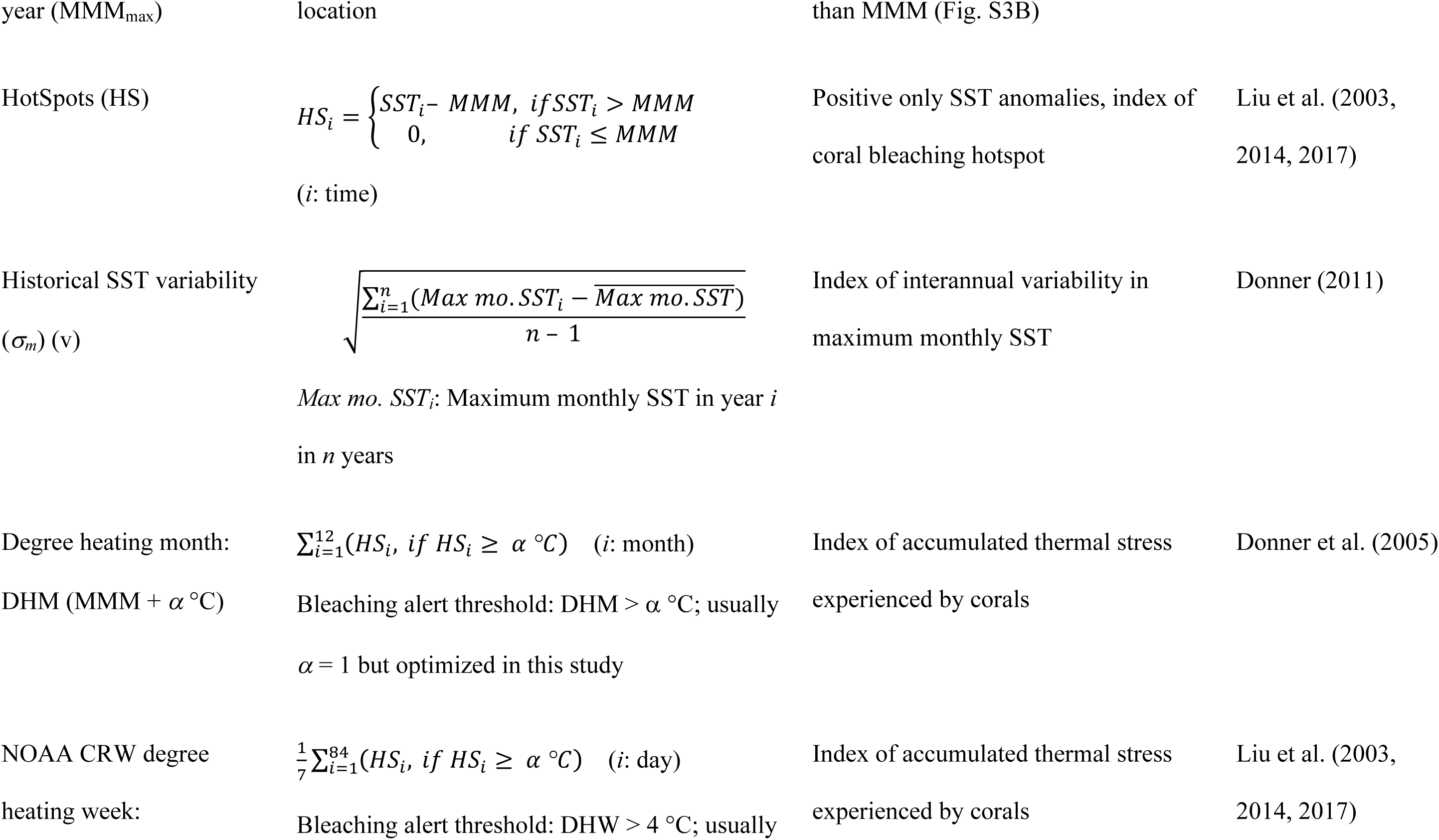
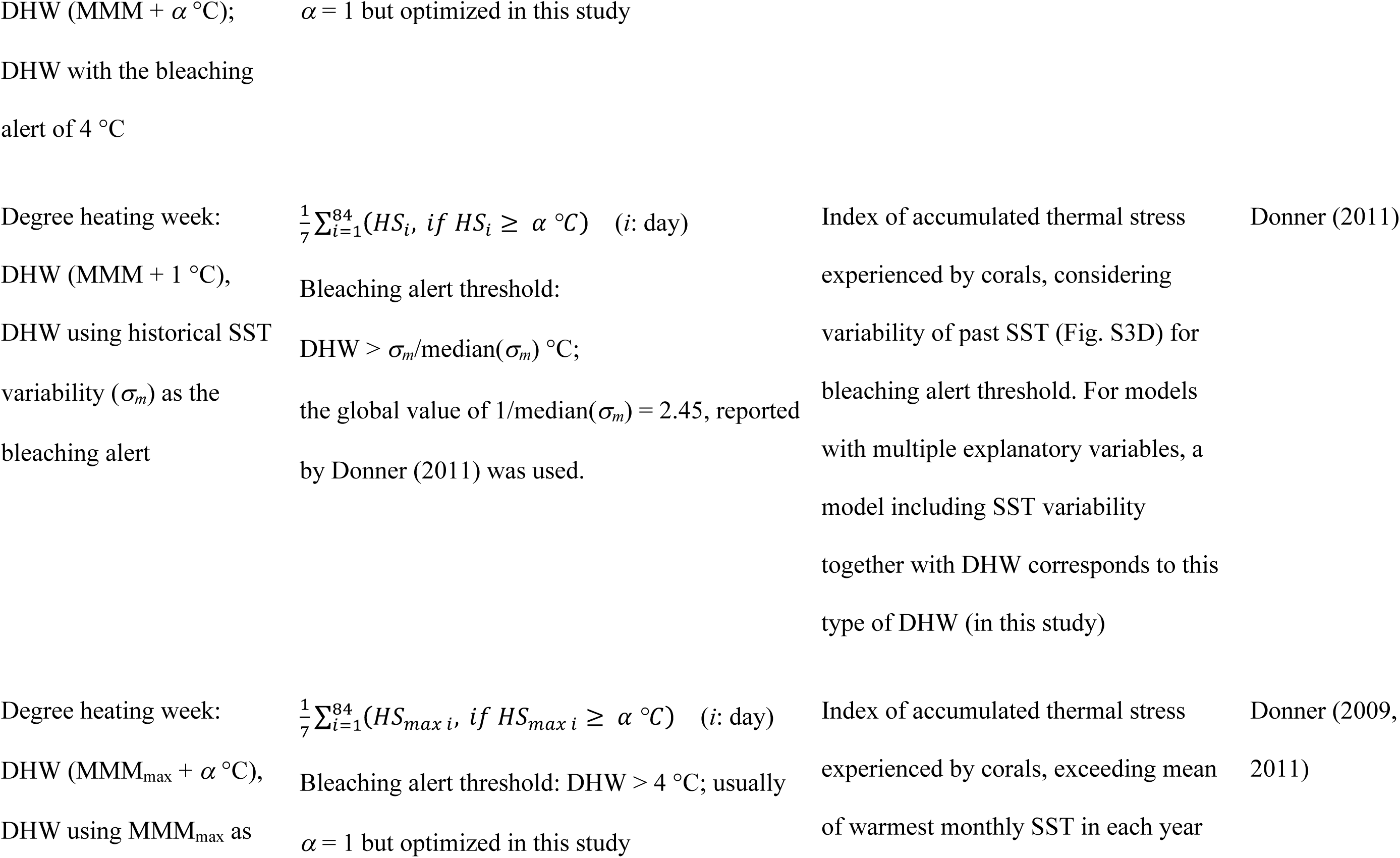
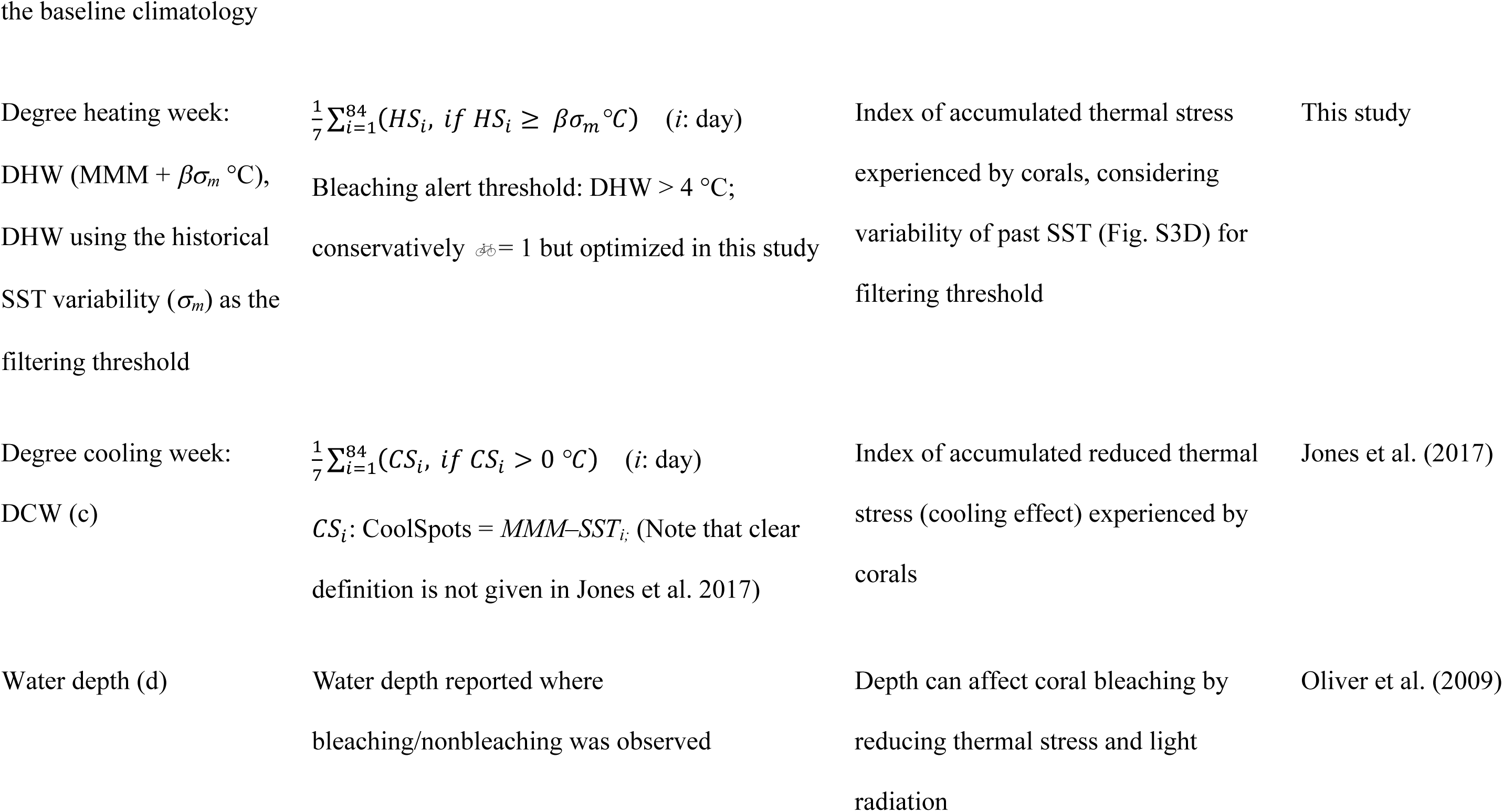
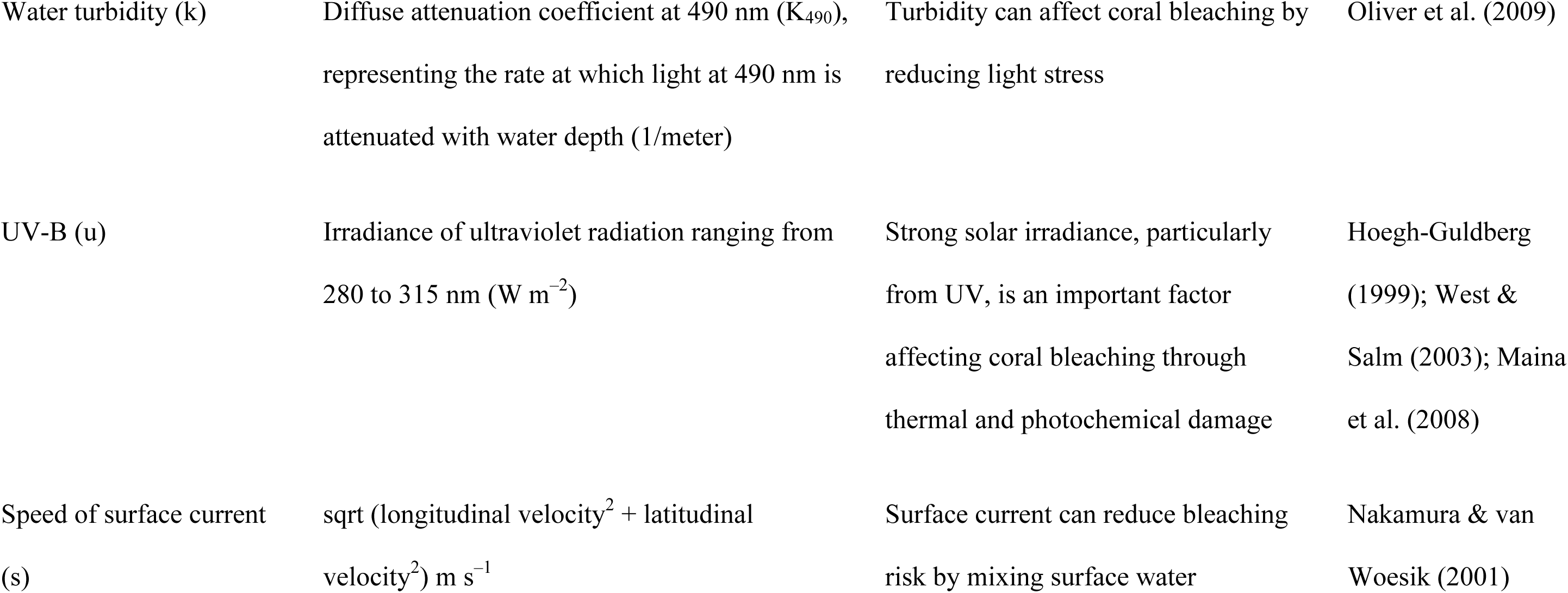
Summary of indices and methods used in this study

## Materials and methods

### Observational records of coral bleaching

We used data derived from the Sango Map Project (http://www.sangomap.jp/) that were submitted up to March 2017. An observer who attempted to submit an observational record to the Sango Map Project was requested to provide the following information as mandatory fields for quality control: (1) his or her handle name and e-mail to identify the observer; (2) presence or absence of corals; (3) the location of the observation according to latitude and longitude, which can be searched through Google Maps API (https://developers.google.com/maps/); (4) the name of the location of the observation; (5) the date, month, and year of the observation; (6) the method of survey (scuba diving, snorkeling, glass boat, walking, or other); (7) the depth of the water at which the observation was made in meters; (8) the observer’s working relationship to coral reefs (professional scientist, nonprofit or nongovernmental organization, tourism, or other); (9) the level of severity of coral bleaching (high, medium, low, or nonbleaching/not found, according to the bleaching dataset of ReefBase); (9) the percentage of corals bleached; (10) the colony form of the bleached corals (arborescent, foliose, massive, table, encrusting, or free-living); (11) the presence or absence of mass mortality in other organisms, to show whether there had been an unusual environmental event causing mass mortality including corals; and (12) remarks about the observation. The observers were also asked to submit photographs of the observation. We confirmed or rejected questionable records (e.g., observations on land, in the open ocean, or of doubtful corals) by questioning the observer via e-mail. Further, Committee Sango-Map-Project held several events and workshops to announce and promote the Sango Map Project for participants and potential observers (Namizaki et al., 2013).

After quality control and exclusion of records without information about bleaching, we obtained 668 totally independent filtered records of observations made between July 2004 and October 2016. Fifty-two of these observations were submitted by professional scientists, 152 by nonprofit or nongovernmental organizations, and 134 by tourists. Fifty-nine observations were conducted as part of CoralWatch (http://www.coralwatch.org/) and 63 as part of ReefCheck Japan(http://www.reefcheck.jp/). The records provided good spatial coverage of the coral reefs in Japan (Fig. 1). Many of the records were obtained in the first 3 years after the Sango Map Project launched, including 449 records during 2008–2010. In addition, 82 and 111 records were reported in 2013 and 2016, respectively, when mass bleaching events were observed in Japan (Kayanne, 2017; Kayanne et al., 2017).

**Figure 1.**
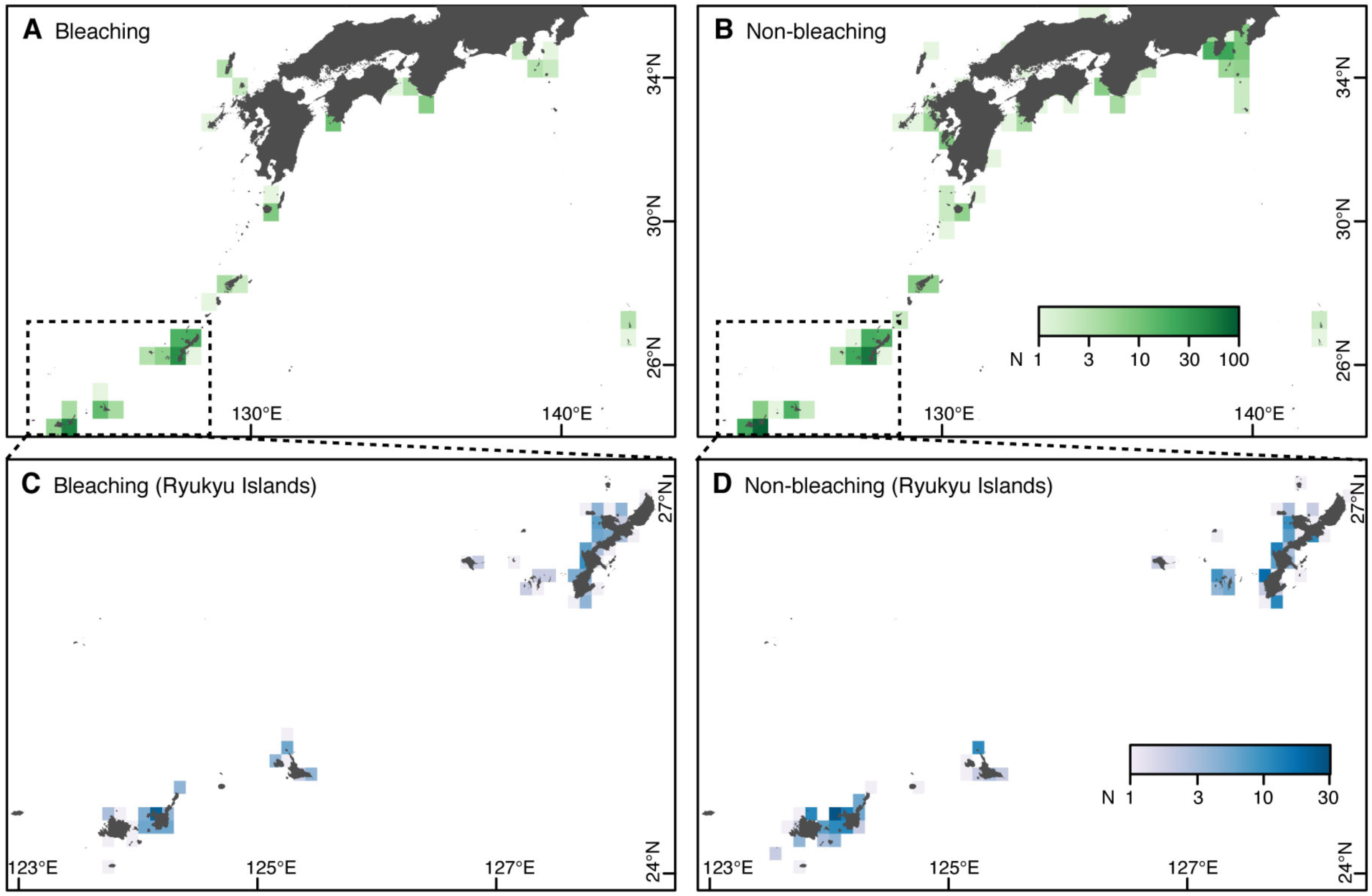
Study area and number of observations in southern Japan. (A, B) Whole study area. (C, D) Main study area, enclosed by a dashed square in A and B. (A, C) Observations of coral bleaching. (B, D) Observations of nonbleaching.

The records included observations of “bleaching” not induced by thermal stress (e.g., small bleaching events within microatolls, disease, or scars by predation) and observations made after the actual occurrence of the bleaching event, e.g., an observer visiting a reef in the autumn might find a coral that had been bleached in the summer. Therefore, such bleaching observations, for which the DHW value at the local 1-km grid on the observation date did not exceed zero, were regarded as nonbleaching observations for our analyses. By this procedure, we reclassified 107 bleaching observations as nonbleaching observations.

After the screening, the prevalence of records was not so biased: consisting of 228 bleaching and 440 nonbleaching observations (Fig. 1), and therefore the risk of biased predictions was low (Table 1). The yearly and spatial patterns of bleaching occurrences were consistent with those in previous reports from Japan (Kayanne, 2017; Kayanne et al., 2017). We checked spatial autocorrelation in the residuals from a prediction model (Table 1) of the NOAA CRW DHW using the spatial autocorrelation coefficient (Moran’s *I*) and confirmed that there was no significant autocorrelation in the residuals, indicating no significant spatial bias in the data (Dorman et al., 2007).

### Thermal indices

To calculate thermal indices, we used the daily data of Multi-scale Ultra-high Resolution Sea Surface Temperature (MUR SST) Analysis version 4.1 (JPL MUR MEaSUREs Project, 2015), in which the spatial resolution is 1 km (0.01°). The MUR SST is a blend of SSTs from six different satellites and thus provides high accuracy. However, the MUR SST is available only from 2002 to the present, which is not a sufficient period to calculate maximum monthly mean (MMM) climatology. Therefore, we used the Optimum Interpolation SST (OI SST) version 2 (Reynolds et al., 2007), provided by NOAA ESRL PSD (http://www.esrl.noaa.gov/psd/) from 1985 to 2002, in addition to the MUR SST. To correct the SST bias between MUR SST and OI SST, we added the bias in monthly climatology of 2002 to the present (May 2017) to OI SST after down-scaling to 0.01° using inverse distance weighting interpolation (Tabor & Williams, 2010; Yara et al., 2011).

Using the monthly mean SSTs from 1985 to 2015, we obtained two types of MMM climatologies. The first MMM climatology follows the protocol in NOAA CRW ver. 3 by Liu et al. (2014, 2017), in which the temporal midpoint was recentered to that of the heritage 50 km MMM (1985–1990, 1993) using the approach of Heron et al. (2014) as follows: *SST*_recentered i_ = *SST*_i_ – *slope*_*i*_ × (*T*_*1985–2015*_ – *T*_*1985–1993*_), where *SST*_*i*_ is the SST climatology as obtained above and *SST*_*recentered* *i*_ is the recentered SST climatology at the cell *i*. *slope*_*i*_ represents the linear trend of monthly mean SSTs between the center times of the two time durations (*T*_*1985–2015*_, *T*_*1985–1993*_) at the cell *i*. The down-scaled and recentered MMM was in good agreement with the CRW MMM ver. 3 (Fig. S2). The second one was known as MMM_max_ climatology (Donner, 2009, 2011), which uses the mean of the warmest month of each year instead of the mean of the warmest month during the climatological years in MMM climatology. The warmest month is not always the same, therefore MMM_max_ is larger than MMM (Fig. S3A-C) and represents the actual seasonal peak in SST better than MMM climatology. This method is particularly effective in tropical zones with less seasonality (Donner, 2011).

We calculated eight types of thermal and cooling indices, including mean weekly and monthly SST, DHM, DHW (MMM + α °C), DHW (MMM + α °C) using SST variation as bleaching alert threshold, DHW (MMM_max_ + α°C), DHW (MMM + βσ_µ_ °C), and degree cooling week (DCW) for each grid cell and observation day (Table 2). The historical SST variability (σ_m_) (Table 2), ranging from 0.36 to 0.71, with a median of 0.57 (Fig. S2D), was calculated using the monthly mean SST from 1985 to 2015. Although DCW is calculated by a similar algorithm as DHW, it was not correlated with DHWs, so we included DCW as a covariate. The filtering threshold (α, β) is fixed to 1 in the standard indices, though we searched for the optimum threshold to maximize predictive performance.

### Other environmental variables

We considered satellite-derived data of UV-B (Table 2) and photosynthetically active radiation (PAR). Monthly data for UV-B and PAR were obtained from Japan Aerospace eXploration Agency Satellite Monitoring for Environmental Studies (JASMES) (http://kuroshio.eorc.jaxa.jp/JASMES/; accessed 25 June 2017), taking the average of MODIS-Aqua and Terra (http://modis.gsfc.nasa.gov/data/dataprod/). Although both UV and PAR have the potential to affect coral bleaching (Hoegh-Guldberg, 1999), they showed strong correlation (*r* = 0.79) as in Yee et al. (2008), i.e., multicollinearity (see Table 1). Therefore, we excluded PAR from our analysis.

Increasing the speed of surface currents and wind can reduce bleaching risk by mixing surface seawater (Nakamura & van Woesik, 2001; Maina et al., 2008). For speed of the surface current, we extracted estimates from the HYCOM+NCODA Global 1/12° Analysis GLBu0.08 for 1997 to 2017 (https://hycom.org/dataserver/gofs-3pt0/analysis/; accessed 27 December 2016). Then, we obtained the climatologic data from July to September, during which most of the bleaching events were observed in this study. As an index of wind speed, typhoon tracking data were obtained from the Regional Specialized Meteorological Center Tokyo (http://www.jma.go.jp/jma/jma-eng/jma-center/rsmc-hp-pub-eg/trackarchives.html; accessed 22 June 2017). We used the length of time without typhoon, defined as a wind speed over 15 m sec^−1^, for each grid cell. However, the length of time without typhoon was strongly correlated with DHW (*r* = 0.86), and therefore we excluded it from our analysis.

We also used the diffuse attenuation coefficient (K_490_) as an index of water turbidity, which can affect coral bleaching by reducing the stress of light radiation (Table 2). A monthly composite of K_490_ (4 km, Level-3 binned MODIS AQUA products) was obtained from the NOAA OceanColor database (http://oceancolor.gsfc.nasa.gov; accessed 8 September 2017).

We obtained the climatology during July to September, as for current speed. Sea floor slope was calculated from the Shuttle Radar Topography Mission digital elevation model (SRTM30; 0.0083 degree) (https://www2.jpl.nasa.gov/srtm/; accessed 28 April 2016).

Data on current speed and diffuse attenuation were down-scaled to 1 km, using bilinear interpolation. If values of environmental variables were not available for coastal cells, we used inverse distance weighting interpolation to estimate them.

### Model evaluation and optimization

We evaluated coral bleaching models based on the accuracy of both positive (bleaching) and negative (nonbleaching) predictions. Many previous studies evaluated models of coral bleaching based only on overall accuracy, such as “proportion of correct predictions” and AIC (e.g., Maina et al., 2008, 2011; McClanahan et al., 2015; Kayanne, 2017; Welle et al., 2017), whereas some studies differentiated between the accuracy of positive and negative predictions (Yee et al., 2008; van Hooidonk & Huber, 2009; Donner, 2011). Under unequal prevalence (e.g., much fewer or more nonbleaching than bleaching observations), this can cause biased predictions. For the purpose of comparison, we used four types of evaluation methods: overall accuracy, true positive rate (TPR), true negative rate (TNR), and true skill statistics (TSS) (Table 2). TSS can be used to represent prediction skill, because the index equally assesses positive and negative predictions (Allouche et al., 2006).

To assess the combined effects of multiple environmental influences in addition to thermal stress on coral bleaching, we also constructed prediction models of bleaching using two modeling methods: generalized linear model (GLM) with a binomial error distribution and a logit link function, and random forest (RF) (Breiman, 2001). Although both models give prediction in the form of probability, the underling algorithms are quite different between the models. GLM is an extension of regression models, whereas RF is a machine learning method using randomly repeating classifications (Table 2). Therefore, the fitted model of GLM can be written as a formula that is easy to be used subsequently. In contrast, RF can only be saved electronically, despite its high predictive performance. We confirmed that the data met the assumption of binomial GLM that the residual deviance per degree of freedom was less than 1.5 (Zuur et al., 2009). RF was performed using the randomForest function of the randomForest R package. RF was used under the standard settings to avoid overfitting to training data, except that we followed the recommendation of Hijmans et al. (2017), using “regression model” even though the response variable was classification. Additionally, the relative importance of variables was calculated using the “importance” function of the MuMIn package (Bartoń, 2015) for GLM and the “importance” function of the randomForest package for RF (Liaw & Wiener, 2002).

The predicted probability was transformed into bleaching and nonbleaching by the threshold that maximizes the sum of TPR and TNR (Liu et al., 2005). As the threshold, 0.5 (i.e., the midpoint) is frequently used, although its transformed prediction of occurrences and absences will be biased, particularly if numbers are unequal between observed occurrences and absences in the source data (Liu et al., 2005). This problem is known in studies of species distribution modeling, although only a few studies of coral bleaching have addressed it (van Hooidonk & Huber, 2009). We used the “evaluation” function of the dismo R package (Hijmans et al., 2017) for model evaluation and optimization of the evaluation threshold.

The models were evaluated by 10 repeated cross-validations, using TSS as the evaluation index. In each repeat, we separated 30% of the data as testing data and used the remaining 70% of the data for constructing GLM and RF (Table 1). We optimized the filtering thresholds for DHM and DHWs by cross-validation, while the filtering threshold is fixed at 1.0 °C in the standard indices (Table 1). We searched the optimum filtering threshold between 0 and 1.5 °C for indices using constant threshold (α), whereas we examined the coefficient of σ_*m*_ (*β*) between 0.1 and 2.5 (Table 1) at 0.01 precision (i.e., 151 and 241 submodels, respectively). For models with multiple explanatory variables, we considered DCW, historical SST variability, UV-B, water turbidity, water depth, and current speed, in addition to a thermal index. The optimum set of explanatory variables was also specified through cross-validation, i.e., selecting the set of variables that best explains the testing data among all 15 possible combinations, excluding the two most influential variables (DCW and UV-B), which are always included in models with multiple explanatory variables (Table 1). Therefore, we totally evaluated 22,650 or 36,150 models (10 cross-validations × 15 variable combinations × 151 or 241 submodels) for each GLM and RF model.

Finally, we predicted coral bleaching in the warmest month of the main coral habitat in the study area using the optimized best prediction model. We also assessed the reduced UV-B effect on coral bleaching as a possible adaptive measure, applying the 40% reduction in UV-B radiation from the original values by the 40% increase in screening effect (water turbidity). These levels of reduction and increase in variables were consistent with *in situ* examination in Onna Village in the Ryukyu Islands (Okinawa Prefecture, 2017). Prediction was performed using each model built in each of the 10 cross-validations and subsequently averaged among the 10 models. The source of the data for the Japanese map was the Global Map Japan version 2.1 Vector data, provided by the Geospatial Information Authority of Japan (2015). All analytical codes (available in Supplemental Information 1) were written in R language and conducted using R ver. 3.4.1 (R Core Team, 2017).

## Results

### Effect of each environmental variable

GLMs using a single environmental variable indicated various relationships with coral bleaching (Fig. 2). The predicted probability of bleaching increased with the thermal indices, including SSTs, DHM, DHWs, and UV-B, and decreased with DCW, water turbidity, and water depth. The responses to historical SST variability and current speed were not significant, with 95% confidence intervals (CIs) ranging from negative to positive responses. Both of the predicted responses to monthly and weekly SSTs were positive, although their 95% CIs were too wide, suggesting they were not reliable indices of coral bleaching. At the same time, the predicted bleaching alert thresholds to discriminate coral bleaching were found to be lower than the standard thresholds, except for DHM (Table 3A).

**Figure 2.**
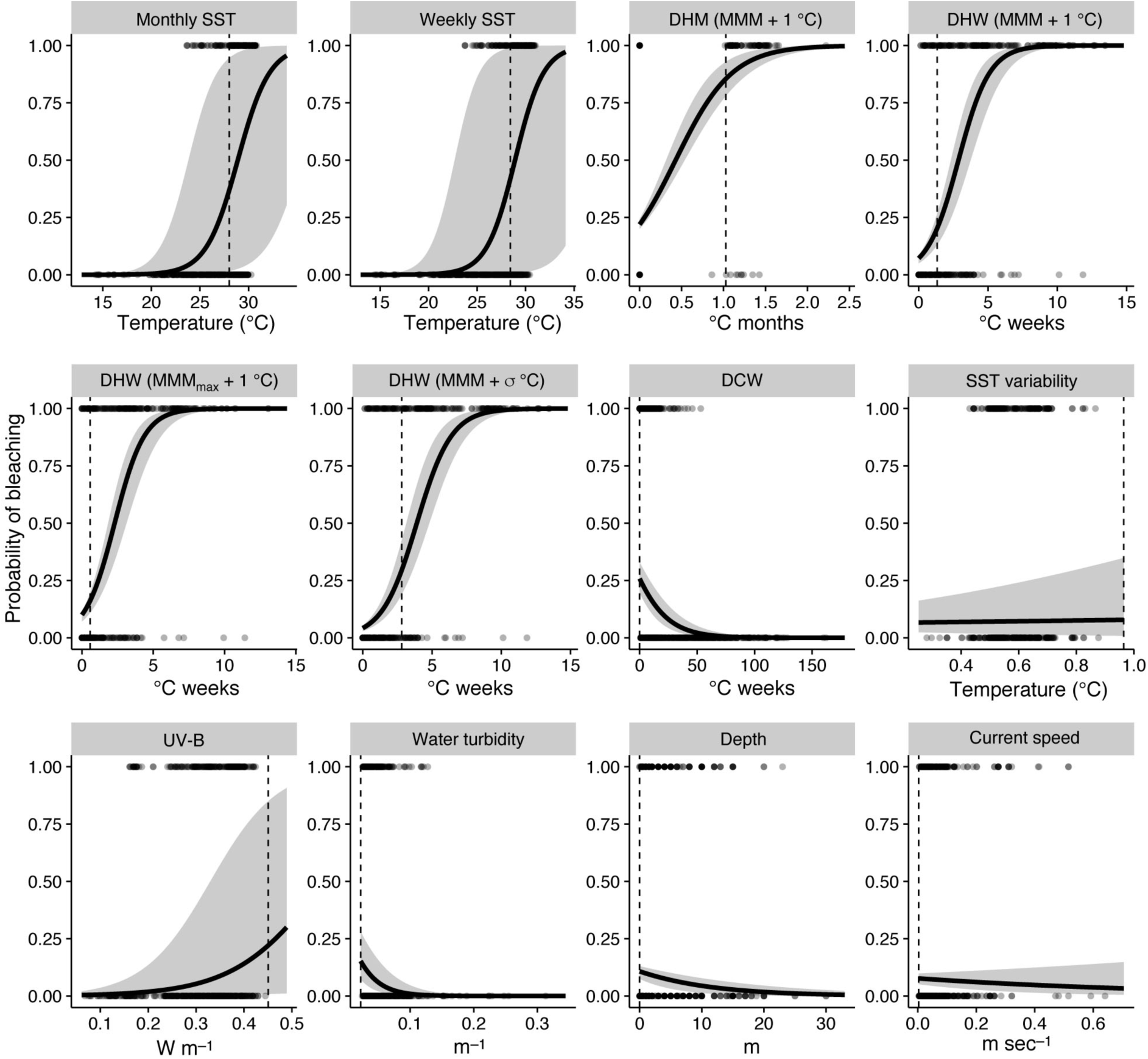
Observed and predicted coral bleaching in relation to each environmental variable. Generalized linear model of each single variable. Values of 1 and 0 represent bleaching and nonbleaching, respectively. Solid lines and gray areas indicate mean model fit and 95% confidential interval, respectively. Dotted line represents the threshold discriminating bleaching and nonbleaching, which was optimized by the true positive rate-true negative rate (TPR-TNR) sum maximization approach (see Table 2).

**Table 3.** Prediction models of coral bleaching with optimized evaluation and filtering thresholds. (A) Thermal index only model with optimized evaluation threshold. (B) Thermal index only model with optimized evaluation and filtering thresholds. (C) Multiple explanatory variables with optimized evaluation threshold and standard filtering threshold. (D) Multiple explanatory variables with optimized evaluation and filtering thresholds. In (C) and (D), c: DCW; d: depth; k: water turbidity; u: UV-B radiation; s: current speed; v: historical SST variability (see Table 2). The optimized evaluation thresholds (mean ± SE) of the predicted probability of coral bleaching are shown with corresponding bleaching alert thresholds of thermal indices. The optimized formula for predicted probability of bleaching is shown for GLM. logistic(x) = 1/ (1 + exp(–x)).

| Model | Evaluation threshold (Bleaching alert threshold °C) | Predicted formula (for GLMs) |
| --- | --- | --- |
| Monthly SST (GLM) | $0.377 \pm 0.010$ (28.01 °C) | $\text{logistic}(-17.7 + 0.612 \cdot \text{SST})$ |
| Monthly SST (RF) | $0.346 \pm 0.010$ | |
| Weekly SST (GLM) | $0.409 \pm 0.012$ (28.04 °C) | $\text{logistic}(-19.7 + 0.68 \cdot \text{SST})$ |
| Weekly SST (RF) | $0.309 \pm 0.020$ | |
| DHM (MMM + 1 °C) (GLM) | $0.855 \pm 0.004$ (1.02 °C) | $\text{logistic}(-1.27 + 2.96 \cdot \text{DHM})$ |
| DHM (MMM + 1 °C) (RF) | $0.454 \pm 0.015$ | |
| DHW (MMM + 1 °C) (GLM) | $0.208 \pm 0.012$ (1.33 °C) | $\text{logistic}(-2.56 + 0.891 \cdot \text{DHW})$ |
| DHW (MMM + 1 °C) (RF) | $0.129 \pm 0.019$ | |
| DHW (MMM<sub>max</sub> + 1 °C) (GLM) | $0.162 \pm 0.009$ (0.58 °C) | $\text{logistic}(-2.2 + 0.958 \cdot \text{DHW})$ |
| DHW ($\text{MMM}_{\text{max}} + 1 \text{ }^{\circ}\text{C}$) (RF) | $0.268 \pm 0.017$ | |
| DHW ($\text{MMM} + \sigma_m \text{ }^{\circ}\text{C}$) (GLM) | $0.292 \pm 0.022$ (2.81 $^{\circ}\text{C}$) | $\text{logistic}(-3.12 + 0.800 \cdot \text{DHW})$ |
| DHW ($\text{MMM} + \sigma_m \text{ }^{\circ}\text{C}$) (RF) | $0.196 \pm 0.017$ | |

(B) Thermal index only model with optimized evaluation and filtering thresholds
| Model | Evaluation threshold<br>(Bleaching alert threshold $^{\circ}\text{C}$) | Filtering threshold | Predicted formula (for GLMs) |
| --- | --- | --- | --- |
| DHM ($\text{MMM} + \alpha \text{ }^{\circ}\text{C}$) (GLM) | $0.464 \pm 0.024$ (0.611 $^{\circ}\text{C}$) | $\alpha = 0.23$ | $\text{logistic}(-1.65 + 2.48 \cdot \text{DHM})$ |
| DHM ($\text{MMM} + \alpha \text{ }^{\circ}\text{C}$) (RF) | $0.369 \pm 0.010$ | $\alpha = 0.23$ | |
| DHW ($\text{MMM} + \alpha \text{ }^{\circ}\text{C}$) (GLM) | $0.213 \pm 0.010$ (2.07 $^{\circ}\text{C}$) | $\alpha = 0.68$ | $\text{logistic}(-3.00 + 0.803 \cdot \text{DHW})$ |
| DHW ($\text{MMM} + \alpha \text{ }^{\circ}\text{C}$) (RF) | $0.207 \pm 0.019$ | $\alpha = 0.52$ | |
| DHW ($\text{MMM}_{\text{max}} + \alpha \text{ }^{\circ}\text{C}$) (GLM) | $0.228 \pm 0.006$ (0.296 $^{\circ}\text{C}$) | $\alpha = 0.64$ | $\text{logistic}(-2.64 + 0.892 \cdot \text{DHW})$ |
| DHW ($\text{MMM}_{\text{max}} + \alpha \text{ }^{\circ}\text{C}$) (RF) | $0.174 \pm 0.014$ | $\alpha = 0.89$ | |
| DHW ($\text{MMM} + \beta \cdot \sigma_m \text{ }^{\circ}\text{C}$) (GLM) | $0.134 \pm 0.002$ (1.58 $^{\circ}\text{C}$) | $\beta = 2.13$ | $\text{logistic}(-2.22 + 1.01 \cdot \text{DHW})$ |
| DHW ($\text{MMM} + \beta \cdot \sigma_m \text{ }^{\circ}\text{C}$) (RF) | $0.146 \pm 0.014$ | $\beta = 1.68$ | |

(C) Multiple explanatories with optimized evaluation threshold and the standard filtering threshold
| Model | Evaluation threshold | Optimized explanatory variables / predicted formula for GLMs |
| --- | --- | --- |
| Non thermal (GLM) | $0.389 \pm 0.011$ | $\text{logistic}(-1.37 - 0.112 \cdot \text{c} - 0.0341 \cdot \text{d} + 3.77 \cdot \text{s} + 7.95 \cdot \text{u})$ |
| Non thermal (RF) | $0.333 \pm 0.008$ | c, k, u, v |
| Monthly SST (GLM) | $0.387 \pm 0.011$ | $\text{logistic}(-12.0 + 0.458 \cdot \text{SST} - 0.094 \cdot \text{c} + 4.10 \cdot \text{s} - 0.722 \cdot \text{u})$ |
| Monthly SST (RF) | $0.311 \pm 0.008$ | SST, c, k, s, u, v |
| Weekly SST (GLM) | $0.348 \pm 0.010$ | $\text{logistic}(-17.2 + 0.667 \cdot \text{SST} - 0.084 \cdot \text{c} + 4.69 \cdot \text{s} - 3.56 \cdot \text{u})$ |
| Weekly SST (RF) | $0.322 \pm 0.005$ | SST, c, k, u, v |
| DHM (MMM + 1 °C) (GLM) | 0.387 ± 0.010 | logistic(1.31 + 3.02·DHM – 0.126·c – 0.026·d + 3.55·s + 2.99·u – 2.69·v) |
| DHM (MMM + 1 °C) (RF) | 0.380 ± 0.009 | DHM, c, k, s, u, v |
| DHW (MMM + 1 °C) (GLM) | 0.326 ± 0.005 | logistic(–1.85 + 0.723·DHW – 0.053·c – 17.4·k + 2.29·s + 10.1·u – 3.57·v) |
| DHW (MMM + 1 °C) (RF) | 0.365 ± 0.008 | DHW, c, k, s, u, v |
| DHW (MMM<sub>max</sub> + 1 °C) (GLM) | 0.325 ± 0.006 | logistic(–3.75 + 0.805·DHW – 0.065·c – 17.7·k + 1.18·s + 11.2·u) |
| DHW (MMM<sub>max</sub> + 1 °C) (RF) | 0.395 ± 0.007 | DHW, c, k, u, v |
| DHW (MMM + $\sigma_m$ °C) (GLM) | 0.323 ± 0.010 | logistic(–1.99 + 0.688·DHW – 0.031·c – 18.3·k + 7.41·u – 3.11·v) |
| DHW (MMM + $\sigma_m$ °C) (RF) | 0.393 ± 0.008 | DHW, c, k, s, u, v |

(D) Multiple explanatories with the optimized evaluation and filtering thresholds
| Model | Evaluation threshold | Filtering threshold | Optimized explanatory variables / predicted formula for GLMs |
| --- | --- | --- | --- |
| DHM (MMM + $\alpha$ °C) (GLM) | 0.388 ± 0.006 | $\alpha = 0.73$ | logistic(1.53 + 2.47·DHM – 0.125·c + 3.47·s + 2.37·u – 3.17·v) |
| DHM (MMM + $\alpha$ °C) (RF) | 0.380 ± 0.009 | $\alpha = 0.02$ | DHM, c, k, u, v |
| DHW (MMM + $\alpha$ °C) (GLM) | 0.354 ± 0.007 | $\alpha = 0.90$ | logistic(–1.99 + 0.717·DHW – 0.048·c – 17.3·k + 2.39·s + 9.99·u – 3.74·v) |
| DHW (MMM + $\alpha$ °C) (RF) | 0.378 ± 0.010 | $\alpha = 0.97$ | DHW, c, k, s, u, v |
| DHW (MMM<sub>max</sub> + $\alpha$ °C) (GLM) | 0.320 ± 0.008 | $\alpha = 0.94$ | logistic(–3.73 + 0.789·DHW – 0.062·c – 19.0·k + 8.77·u) |
| DHW (MMM<sub>max</sub> + $\alpha$ °C) (RF) | 0.400 ± 0.008 | $\alpha = 0.87$ | DHW, c, u, v |
| DHW (MMM + $\beta \cdot \sigma_m$ °C) (GLM) | 0.336 ± 0.005 | $\beta = 1.83$ | logistic(–3.15 + 0.773·DHW – 0.053·c – 19.2·k + 8.77·u) |
| DHW (MMM + $\beta \cdot \sigma_m$ °C) (RF) | 0.394 ± 0.006 | $\beta = 1.67$ | DHW, c, d, k, s, u, v |

We compared the importance of different environmental variables (Fig. 3). The ranking of variables was almost the same for GLM and RF: the best was DHW, followed by DCW. UV-B, water turbidity, and historical SST variability were also found to be good explanatory variables for coral bleaching. Historical SST variability and current speed were not the worst variables, despite their inconsistent relationships with coral bleaching, as shown above.

**Figure 3.**
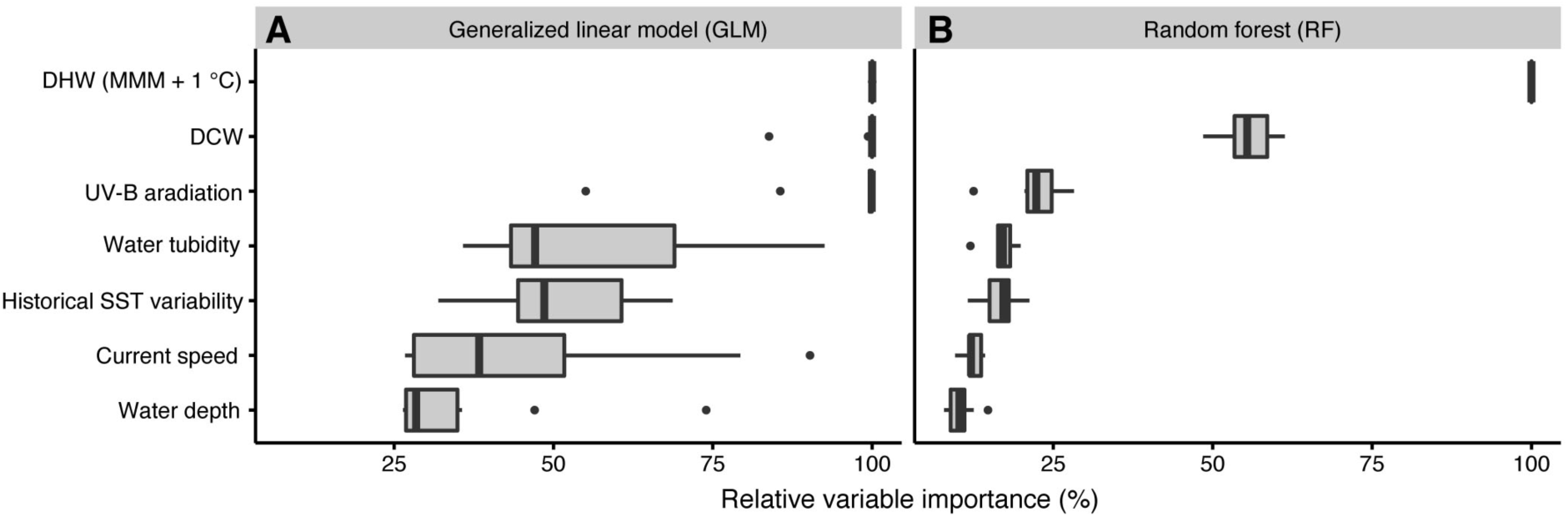
Relative importance of each environmental variable. Under (A) generalized linear model (GLM) and (B) random forest (RF). DCW, degree cooling week; DHW, degree heating week; MMM, maximum monthly mean; SST, sea-surface temperature; UV-B, ultraviolet B.

### Optimization and assessment of the filtering threshold

Optimization of the filtering threshold improved the predictive performance of DHM and DHWs, although the improvement was particularly small for models consisting of multiple environmental variables, whereas the improvement was significant for models consisting of thermal index only (Fig. 4).

**Figure 4.**
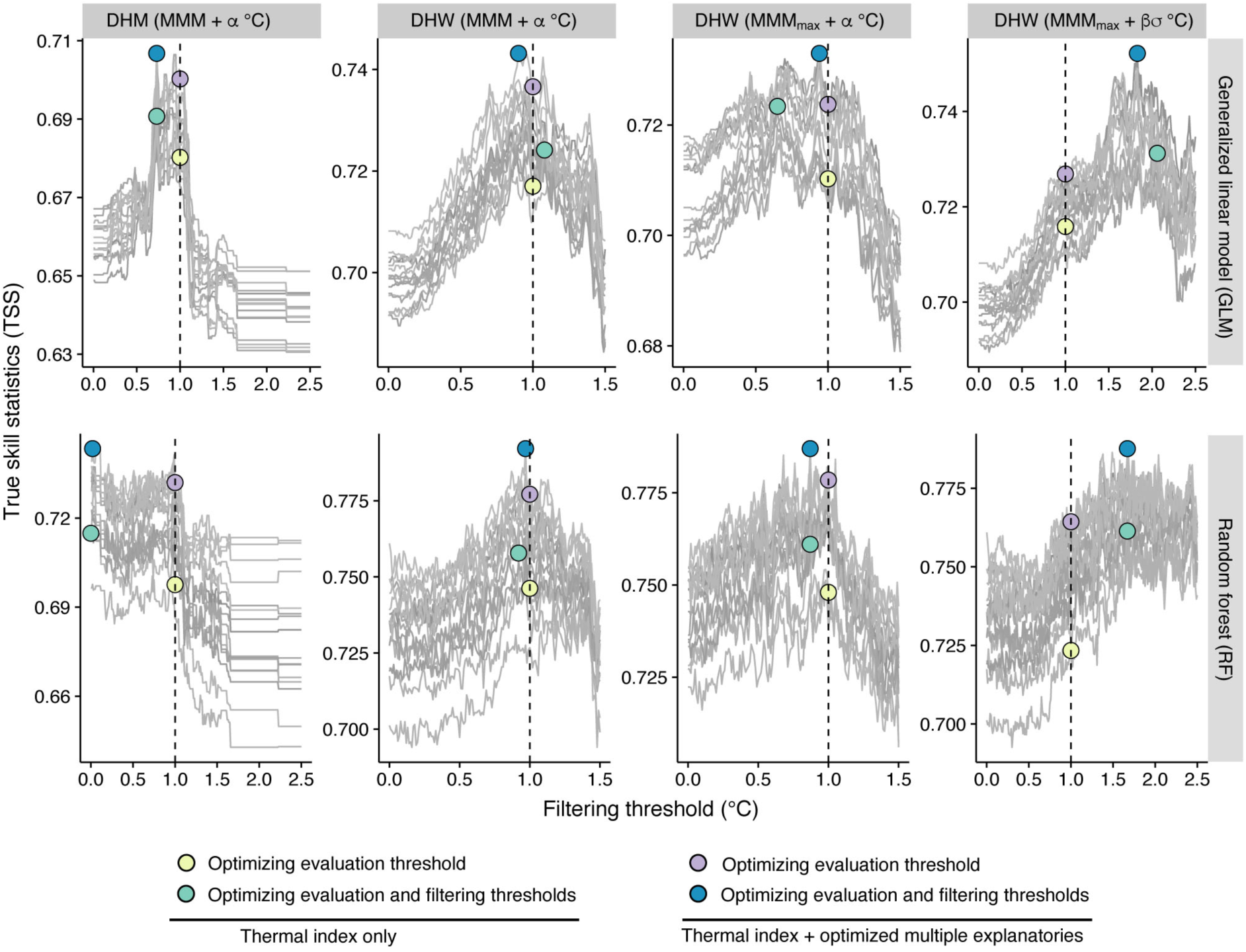
Optimization of filtering threshold. The predictive performance (true skill statistics [TSS]) of each model with the change in filtering threshold for four thermal indices under generalized linear model (GLM) (upper) and random forest (RF) (bottom) (see Tables 1, 2). Individual gray lines represent each of the 15 model combinations of environmental variables. Colors of filled circles correspond to the groups in Figure 5 in relation to optimizing thresholds and explanatory variables. DHM, degree heating month; DHW, degree heating week; MMM, maximum monthly mean.

Further, we compared the predictive performance among all the models including the standard or the optimized thermal index, and the optimized set of multiple explanatory variables (Fig. 5). In thermal index only models with the standard threshold, TNR was larger than TPR, indicating that large overall accuracy can result from large TNR despite small TPR. At the same time, TSS well represented both TPR and TNR. Among the thermal indices with the standard threshold, DHW using historical SST variability (σ_*m*_) as the bleaching alert threshold and that using σ_*m*_ as the filtering threshold equally scored the best performances in TSS (Fig. 5). Weekly SST had no prediction skill (TSS = 0), and the skills of monthly SST were also quite low. TPR of DHW using historical SST variability as the bleaching alert was the largest, although it suffered from the largest false positive rate (1–TNR). Using historical SST variability as the filtering threshold for DHW decreased the false positive rate but also decreased TPR. DHW using MMM_max_ as the baseline climatology showed the worst performance among DHWs, because it had the lowest TPR.

**Figure 5.**
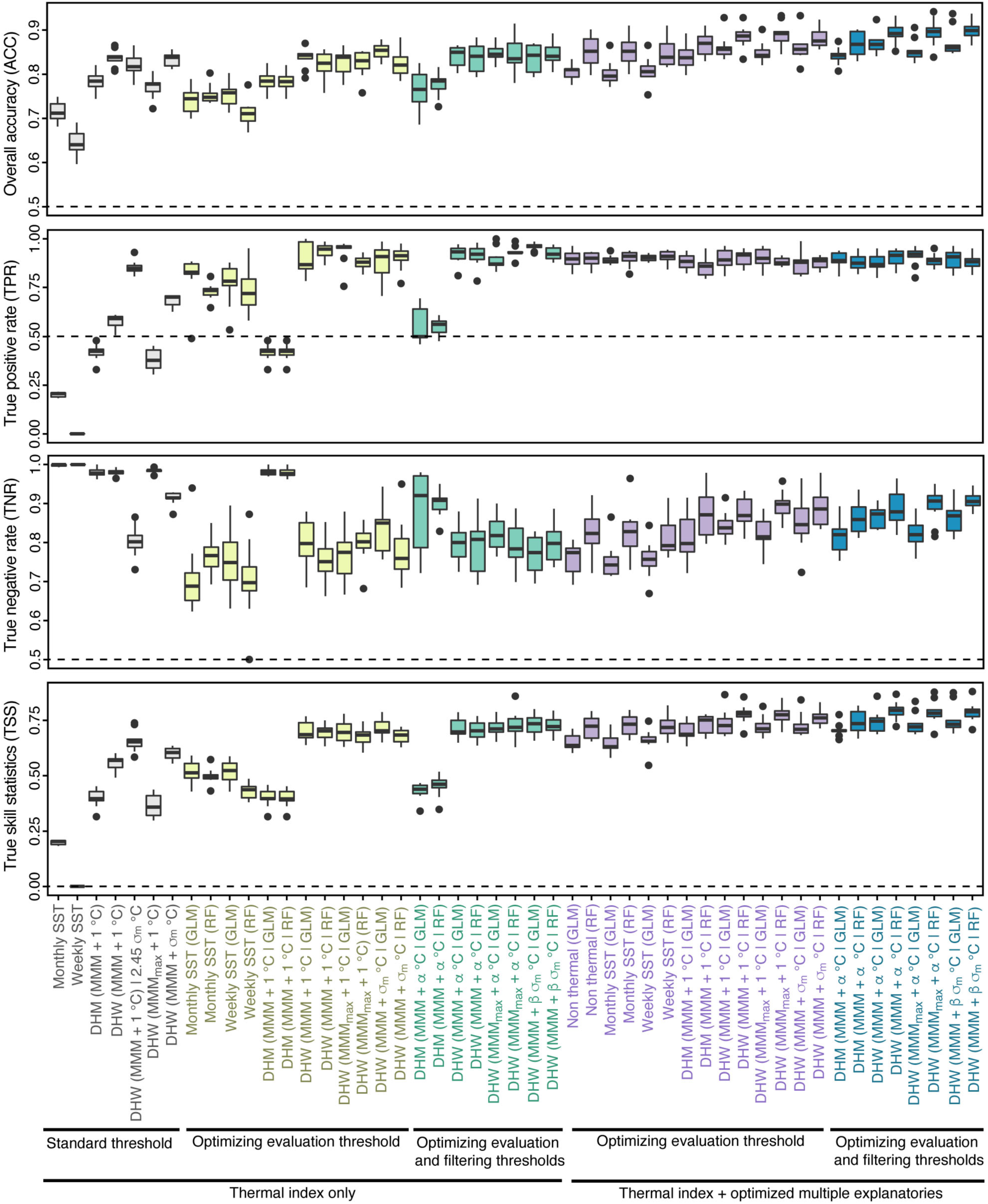
Evaluation of models of coral bleaching. Models are grouped (under bars and colors) according to whether these thresholds are the standard ones or optimized, using thermal index only or thermal index and multiple explanatory variables. The label of a model indicates “index (filtering threshold, if present | abbreviation of statistical model).” E.g., DHW (MMM + β σ_m_ °C | RF) represents the random forest model, including DHW using MMM + β σ_m_ °C as the filtering threshold. See Table 2 for terminology and Table 3 for the optimized evaluation and filtering thresholds and combination of explanatory variables. ACC, overall accuracy; DHM, degree heating month; DHW, degree heating week; GLM, generalized linear model; MMM, maximum monthly mean; RF, random forest; SST, sea-surface temperature; TNR, true negative rate; TPR, true positive rate; TSS, true skill statistics.

Predictive performance was much improved by optimizing the evaluation threshold (Fig. 5, Table 3B). Under the optimized evaluation threshold, DHW with the historical SST variability as the filtering threshold had the best performance among the three types of DHW although the differences were slight, whereas DHM had the worst score. When both the evaluation and the filtering thresholds were optimized (Table 3C), the improvements from the optimizing evaluation threshold were only slight. DHW with the historical SST variability as the filtering threshold still had the best performance under the optimizations. The optimized filtering thresholds were less than 1 °C in all the thermal index only models using the optimized DHM or DHW other than DHWs using σ_*m*_ as the filtering threshold (see also Fig. S3D for σ_*m*_ distribution).

The predictive skill of models with multiple explanatory variables was slightly improved from that of models including only thermal index in the TSS, although the improvement was mainly due to increase in TNR (i.e., reduction of the false positive rate) (Fig. 5). Even models including no thermal index showed high prediction skill. The difference in predictive performance between models with multiple explanatory variables with optimized evaluation thresholds and those with optimized evaluation and filtering thresholds was smaller than that between GLM and RF: RF models were always better than GLM among models with multiple explanatory variables. Although the TPRs of GLMs were better than those of RFs in most cases, the TNRs of GLMs were worse than those of RFs in all models with multiple explanatory variables; i.e., the risk of false positives was higher. Among the models with multiple explanatory variables, the RF model consisting of DHW with MMM + 0.97 °C filtering threshold, DCW, UV-B, water turbidity, historical SST variability, and current speed (Table 3D) had the best prediction skill, with 0.79 for TSS, 0.90 for TPR, and 0.89 for TNR; i.e., it was the best model among all tested. Among the GLMs, the model consisting of DHW with MMM + 1.83·σ_*m*_ °C filtering threshold, DCW, UV-B, and turbidity had the best skill.

### Prediction of coral bleaching

We showed the predicted probability of coral bleaching by the optimized best multivariate model of RF using DHW with MMM + 0.97 °C filtering threshold for the main coral-habitable areas in Japan (Fig. 6). The mean predicted probability of bleaching ranged from 0.46 to 0.74. Spatial variations in the probability of bleaching were found in both the eastern (Fig. 6A) and the western (Fig. 6C) Ryukyu Islands. Some “hotspots” with higher bleaching probabilities were found in the southeastern part of Okinawa Island, the eastern part of the Kerama Islands (Fig. 6A), and the northern part of Ishigaki Island (Fig. 6C). This resulted in bleaching in 2.5 to 5 (44% to 100%) of 5 years (2008–2010, 2013, and 2016) (Fig. 6B, D). Further, we considered the 40% reduction in UV-B radiation by the 40% increase in screening as an adaptation measure against temperature warming (Fig. 7). Under reduced UV-B radiation, the decrease in the predicted probability of bleaching was conspicuous (Fig. 7A, C), particularly in the bleaching hotspots mentioned above (Fig. 6A, C). After UV reduction, the predicted probability of bleaching ranged from 0.34 to 0.63 (Fig. 7A, C), and bleaching frequency ranged from 28% to 92% (Fig. 7B, D). The decrease in probability was up to 0.24, resulting in a significant decrease in bleaching frequency of up to 56% (Fig. 7B, D), with bleaching in fewer than 3 out of 5 years in most areas.

**Figure 6.**
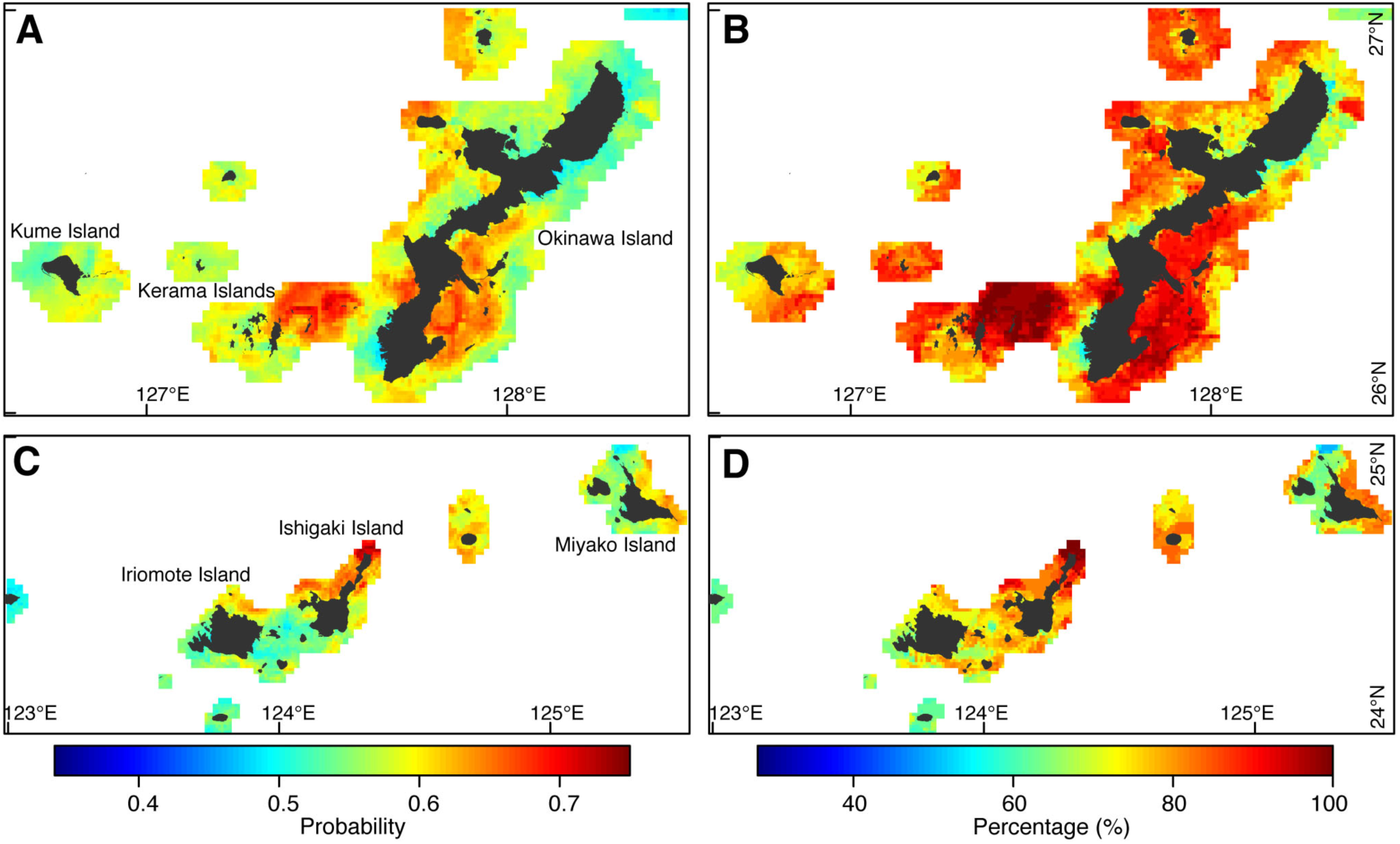
Coral bleaching prediction under the observed environmental settings. (A, B) Eastern Ryukyu Islands. (C, D) Western Ryukyu Islands. (A, C) Mean of the bleaching probabilities in the warmest months among 2008–2010, 2013, and 2016. (B, D) Percentage of predicted bleaching frequencies for 2008–2010, 2013, and 2016; i.e., 100% indicates bleached in all of the years. The average of 10 models built in cross-validations is shown.

**Figure 7.**
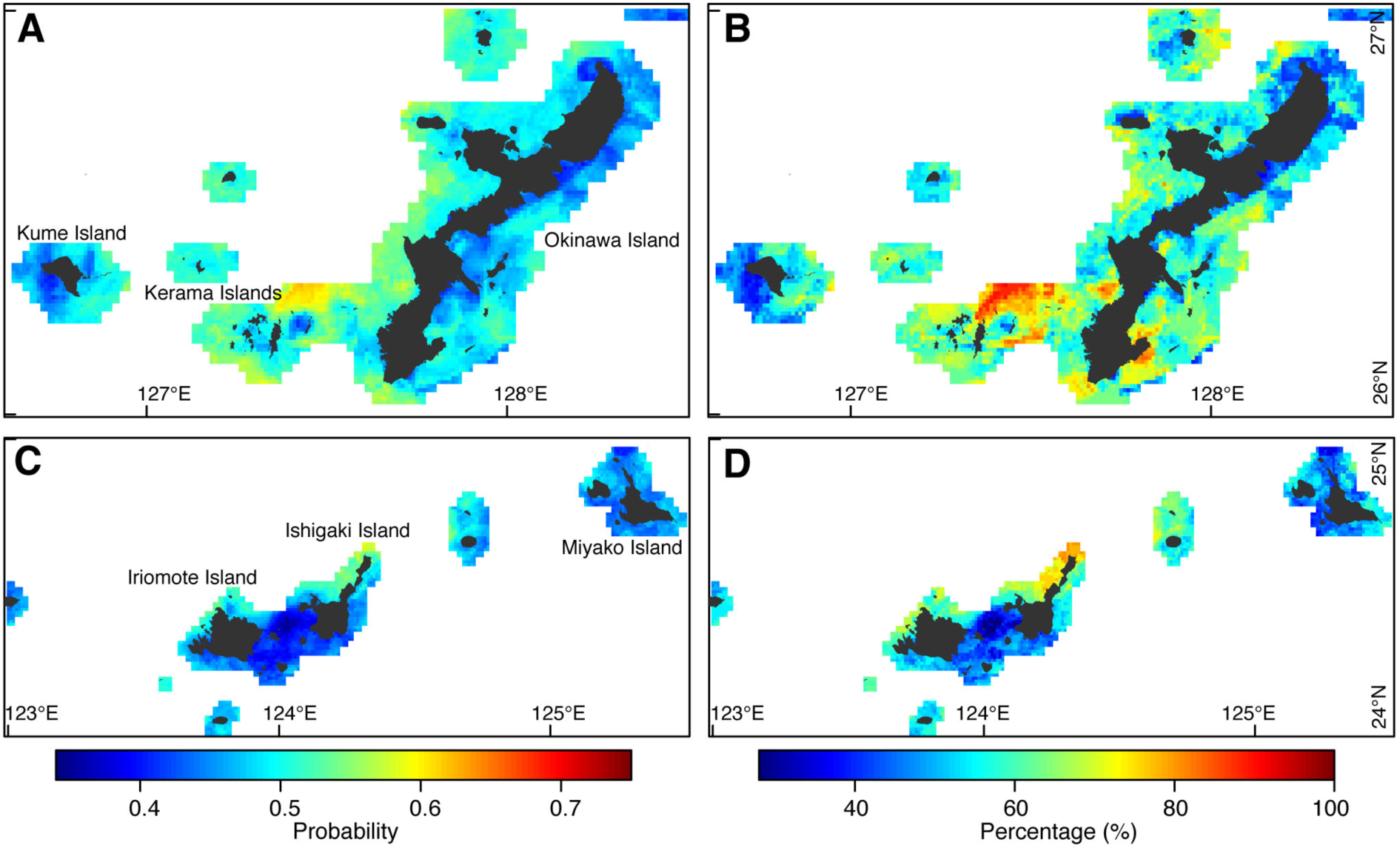
Coral bleaching prediction under reduced ultraviolet B (UV-B) radiation by increased screening effect. Prediction under 40% reduction in UV-B radiation by 40% increase in screening effect. (A, B) Eastern Ryukyu Islands. (C, D) Western Ryukyu Islands. (A, C) Mean of the bleaching probabilities in the warmest months among 2008–2010, 2013, and 2016. (B, D) Percentage of predicted bleaching frequencies for 2008–2010, 2013, and 2016; i.e., 100% indicates bleached in all of the years. The average of 10 models built in cross-validations is shown.

## Discussion

### Optimizing coral bleaching models

By optimizing thermal indices, the ability to predict coral bleaching was much improved, particularly in SST and DHWs. TPRs were mostly lower than TNRs, as in previous studies (e.g., Donner, 2011); however, not only TNR, but also TPR, was sufficiently improved in this study by model optimizations. Although the sole contribution of optimizing the filtering threshold was small, the optimizations, together with evaluation threshold and combination of environmental variables, achieved around 0.9 of TPR and TNR. We also found that cooling (i.e., DCW), UV-B, and screening (i.e., turbidity) effects were good explanatory factors for coral bleaching, particularly under RF models.

The use of various thermal indices, including DHW by NOAA CRW, some other versions of DHWs, and DHM, to predict coral bleaching has been tested. Donner (2011) showed that DHW using historical SST variability as bleaching alert threshold had a higher TPR but a higher false positive rate than NOAA CRW’s DHW, whereas DHW using MMM_max_ had the worst performance but is suitable in equatorial zones, largely in consistency with our results. Because our study was conducted at a higher latitude, the method using MMM_max_ showed the lowest performance among the DHWs, as expected. At the same time, we also utilized historical SST variability as the filtering threshold, and this method had the highest prediction skill through the optimizations among the thermal index only models.

Historical SST variation σ(_*m*_) might be particularly effective in our case, because it was larger in the main area of our study (ca. 0.56) than in Donner’s (2011) study (ca. 0.25). This may be because the Japanese southern islands are encircled by a strong boundary current (Kuroshio) flowing poleward, and tropical water brought by the current can cause significantly faster warming than the global average (Wu et al., 2012).

The optimized filtering thresholds were less than 1 °C in GLMs and RF using only the thermal index. Indeed, many previous studies suggest that even thermal stress not exceeding 1 °C can induce coral bleaching (Brown, 1997; McWilliams et al., 2005; Kleypas et al., 2008). Nevertheless, the threshold had not been statistically optimized before our study. Although RF is new to studies of predicting coral bleaching, we found that RF was an excellent method of predicting coral bleaching.

Models with multiple environmental variables are increasing and are explaining variations in coral bleaching well (Maina et al., 2008; Yee et al., 2008; McClanahan et al., 2015; Welle et al., 2017). Among environmental variables other than thermal indices, UV radiation was a prominent explanatory factor for coral bleaching, in consistency with previous studies (Hoegh-Guldberg, 1999; Maina et al., 2008, 2011; McClanahan et al., 2015). Other variables related to cooling (DCW) (Jones et al., 2017) and screening (water turbidity) (West & Salm, 2003; Oliver et al., 2009; Maina et al., 2011; Oxenford & Vallés, 2016) were also good explanatory factors. Additionally, topographic variables on a smaller scale, including water depth, are known to help reduce thermal stress on corals (West & Salm, 2003; Oliver et al., 2009). Strong winds may also reduce bleaching risk (Maina et al., 2008, 2011; Yee et al., 2008; McClanahan et al., 2015; Welle et al., 2017). This variable was excluded from the tests in our study because of its strong correlation with thermal indices, but it supports previous evidence. There may be some caveats to our prediction, because environmental variations at spatially and temporally small scales have not been sufficiently included. For example, small-scale water flow may improve the resistance of corals to bleaching (Nakamura & van Woesik, 2001), but the 8-km resolution of current speed in our study is too coarse to represent such effects. The microstructure of the sea floor at the meter scale may also be related to local water flow or shading of corals (West & Salm, 2003; Oliver et al., 2009).

In addition to overall bleaching responses, as in our study, thermal threshold also varies among coral species (Maynard et al., 2008; Yee et al., 2008; Guest et al., 2012; Harii et al., 2014; McClanahan, 2014). Branching corals of *Acropora* and *Pocillopora* spp. are more susceptible to thermal stress than massive corals like *Porites* spp. (Maynard et al., 2008; Yee et al., 2008; Guest et al., 2012; Harii et al., 2014; McClanahan, 2014). Further, thermal tolerance increases with repeated excessive thermal stresses (Brown et al., 2002; Maynard et al., 2008; Guest et al., 2012). The potential of corals to adapt to thermal stress has been recognized (Brown et al., 2002; Maynard et al., 2008; Guest et al., 2012), but it was not fully explained statistically in our analysis using historical SST variability and in other previous studies (Donner, 2011; McClanahan et al., 2015). Such variations and changes in thermal tolerance can result from interspecific differences (Maynard et al., 2008; Guest et al., 2012; Harii et al., 2014; McClanahan, 2014) in acclimation, genotypes, and epigenetics of host corals and symbiotic algae (Palumbi et al., 2014; Torda et al., 2017); the effect of such differences is still an open question.

### Coral bleaching in Japan and reef management

Three previous studies (Strong et al., 2002; Harii et al., 2014; Kayanne, 2017; Kayanne et al., 2017) in the Japanese region analyzed coral bleaching occurrences using temperature anomalies and DHW at coarse resolutions of more than 50 km. Coarse DHW catches overall regional trends in the onset of coral bleaching, although it fails to predict bleaching within well-developed reefs (Strong et al., 2002; Harii et al., 2014). Indeed, our bleaching prediction for the Ryukyu Islands using DHW at 1-km resolution improved the TPR to 0.58 from the score of 0.44 (a 14% improvement) calculated from the tables of Kayanne (2017). The improvement resulting from the increase of spatial resolution of SST may not be very much, although the predictive performance is also improved by optimizing thermal thresholds of DHW. Further, prediction of coral bleaches was much improved by incorporating other environmental variables, such as cooling effect, UV radiation, and turbidity, in addition to thermal indices.

The high performance of our bleaching model at 1-km resolution has practical implications for regional and local management of coral reefs. The prediction revealed high frequencies of coral bleaching in many parts of the Ryukyu Islands. Note that spatial variation was found in the predicted bleaching frequency, and the prediction was based on the lowest rank of severity of bleaching, resulting in overemphasis of the bleaching risk.

Practical management to reduce the risk of coral bleaching should include control of coastal water turbidity (Fabricius, 2005). Increase in water turbidity by terrestrial runoff may decrease the resistance (Wooldridge & Done, 2009) or resilience (Hongo & Yamano, 2013) of corals to bleaching. Nevertheless, a recent study showed that turbid coastal regions might be refuges from future climate warming because of their reduced temperature elevation and solar radiation (Cacciapaglia & van Woesik, 2016). However, coastal turbidity has other confounding effects on corals, including increases in coral diseases and space-competing algae (Fabricius, 2005), and therefore coastal turbidity should be carefully managed. Our study showed that reducing UV radiation by increasing screening significantly reduced bleaching risk as an adaptive measure against climate warming. In Onna Village of the Ryukyu Islands, in situ reduction of UV radiation above corals, without increasing water turbidity, has already been tried (Okinawa Prefecture, 2017). In the trial, reduction of UV radiation by 30% to 44% by screening with large fishery nets resulted in a survival rate of 80% in cultured coral colonies in the summer of 2016. In that year, the most severe thermal stress was recorded during the study period from 2004 to 2016 (Kayanne et al., 2017). The reduction in UV radiation was similar to that in our study (40%), and therefore our prediction could provide a quantitative basis for future reef management in this area. We believe that this is an ideal case of a study based on citizen science feeding back a useful outcome to the local community.

## Conclusions

We have shown how the ability to predict coral bleaching is improved by the optimization of thresholds and the use of multiple environmental influences and modeling methods. Both high-resolution modeling and observational records (i.e., the Sango Map Project) enabled high performance of bleaching prediction (Oliver et al., 2009). These results may be useful to other researchers for selecting a prediction method according to their needs or skills. Our high-resolution predictions provide a quantitative basis for regional and local management of reefs (West & Salm, 2003). Although recent Japanese coral reefs are suffering from severe bleaching risks, we show a practical way to reduce the stress by reducing UV radiation. Future studies should address more practical bleaching models at finer spatial resolutions incorporating variations in intrinsic thermal tolerance, historical effects of previous thermal impacts, and local environmental conditions.

## Acknowledgments

This work was supported by the SOUSEI and TOUGOU Program of the Ministry of Education, Culture, Sports, Science, and Technology in Japan (MEXT) and by the Environment Research and Technology Development Fund (S15) of the Ministry of the Environment, Japan. We thank the 244 participants in the Sango Map Project who provided valuable data on coral bleaching events.

## Author contributions

N.H.K and H.Y. conceived the study. H.Y. and C.S. constructed the system of the Sango Map Project. N.H.K. conducted the analysis. N.H.K. drafted the manuscript with substantial input from H.Y. and C.S.

---

**Figure S1.**
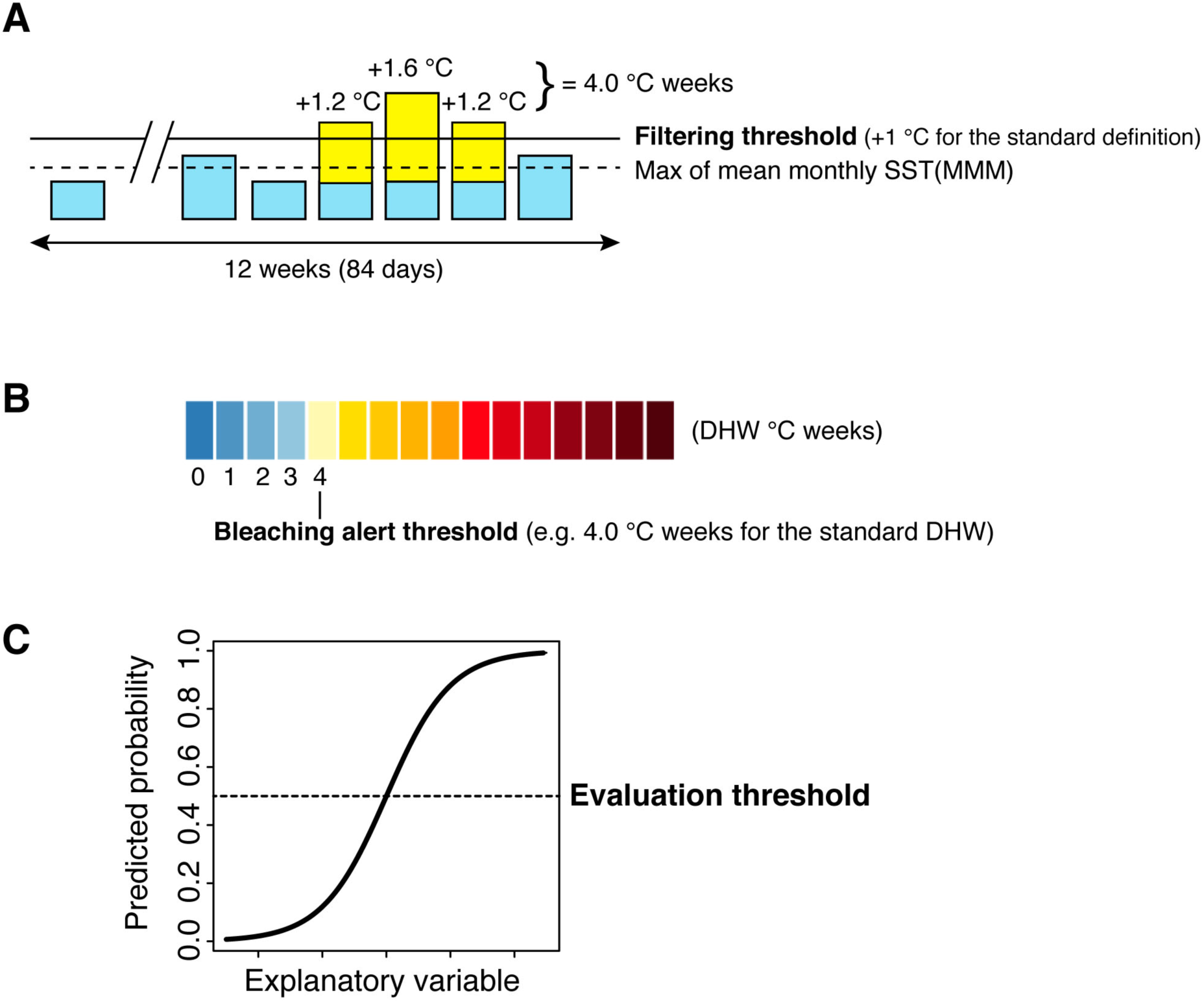
Schematics of thresholds. (A) Filtering threshold: the threshold to filter thermal stress in calculating degree heating weeks (DHWs) or degree heating month (DHM). (B) Bleaching alert threshold: the threshold to discriminate occurrence and absence of bleaching from DHWs or DHM. (C) Evaluation threshold: the threshold to discriminate occurrence and absence of bleaching from the predicted probability of statistical models of bleaching. MMM, maximum monthly mean; SST, sea-surface temperature.

**Figure S2.**
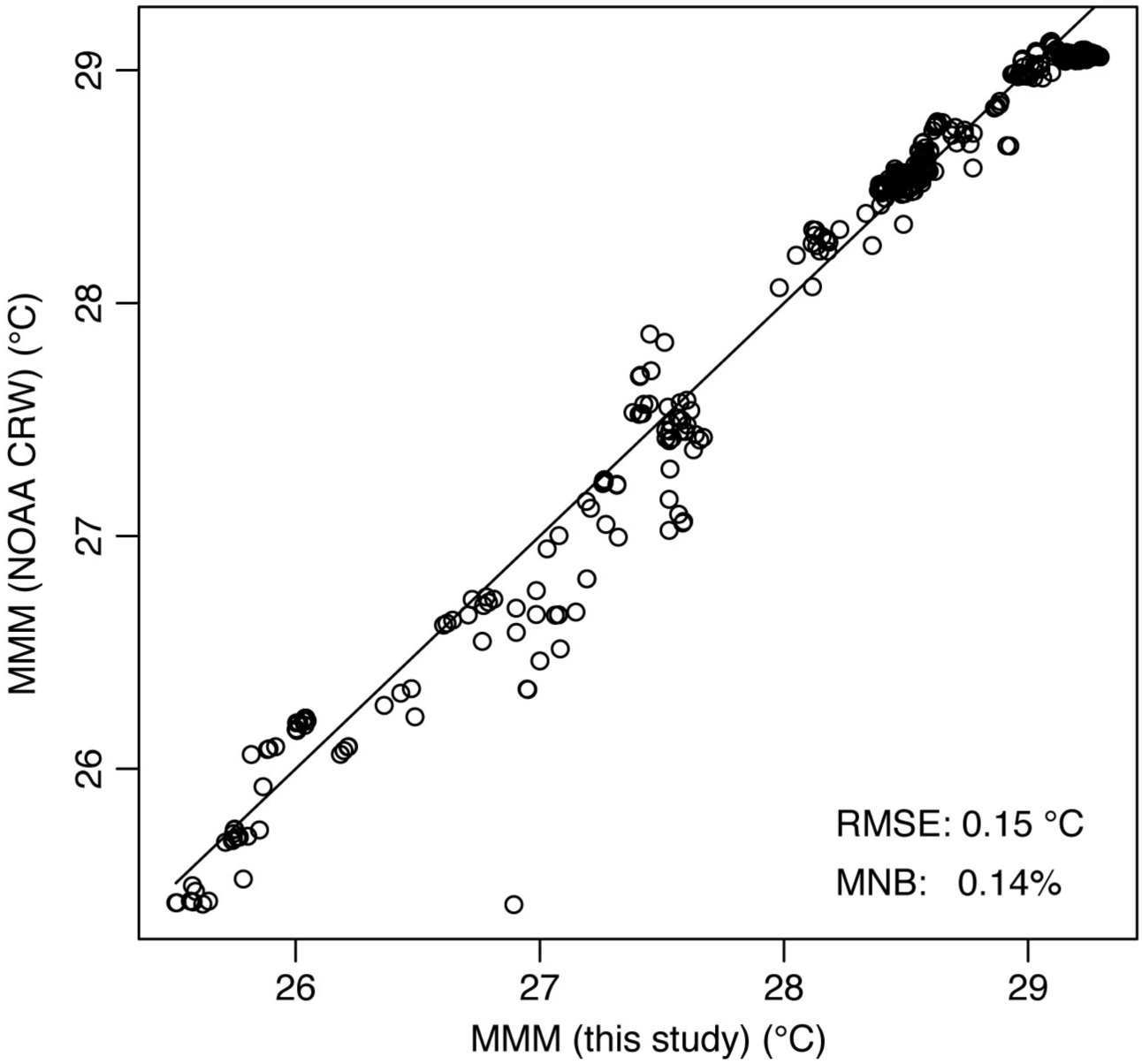
Comparison between MMM SST climatology of NOAA CRW and this study. NOAA CRW MMM was down-scaled to the resolution of MMM of this study using a bilinear interpolation for the comparison. MMM SST, maximum of the monthly mean sea-surface temperature climatology of 1985–2015; MNB, mean normalized bias; NOAA CRW, National Oceanic and Atmospheric Administration Coral Reef Watch; RMSE, root mean square error.

**Figure S3.**
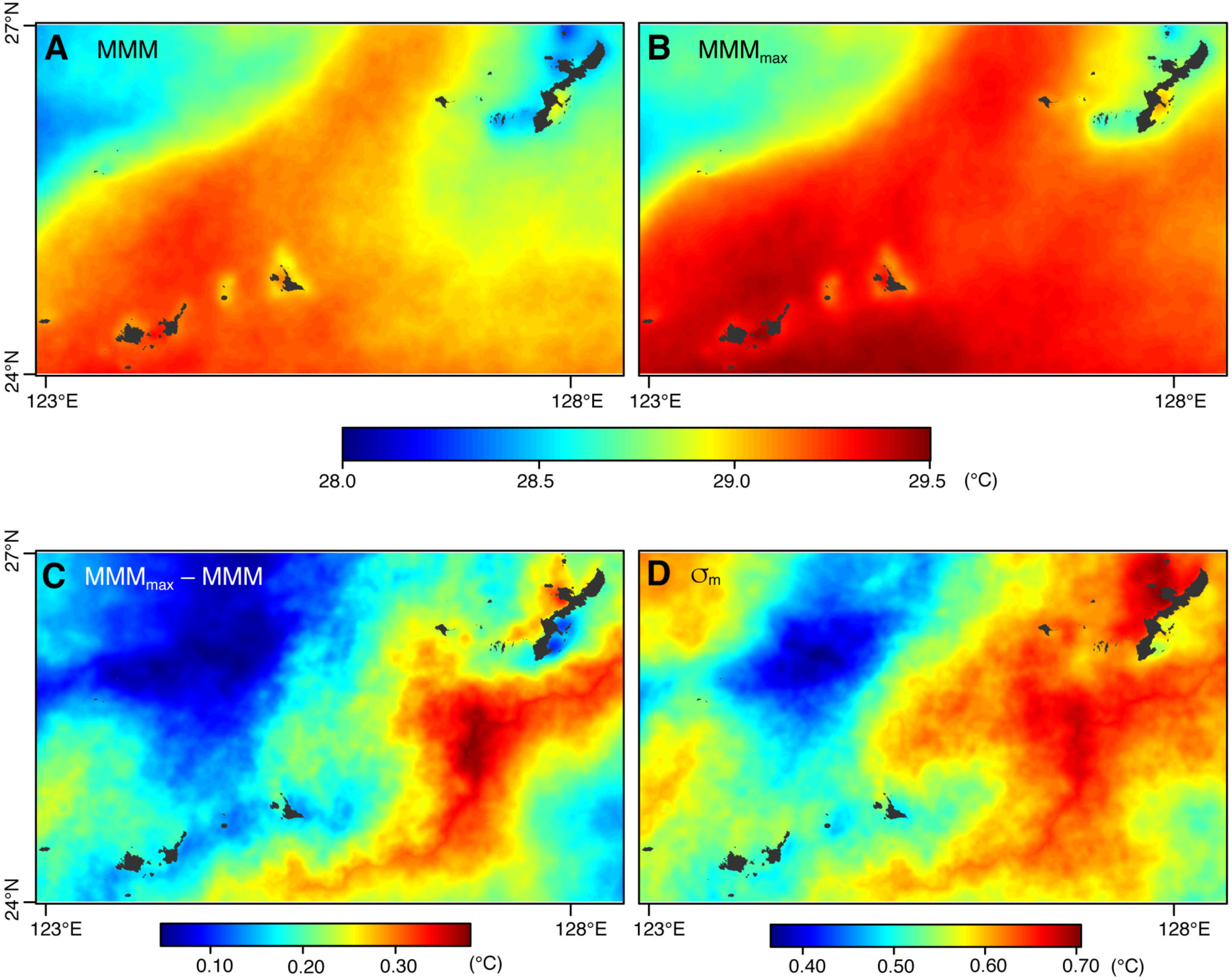
SST climatologies in the Ryukyu Islands (main study area). (A) MMM SST climatology. (B) MMM_max_ SST climatology. (C) Difference between MMM_max_ and MMM. (D) Historical SST variability (standard deviation of maximum monthly SST). MMM, maximum monthly mean; MMM_max_; SST, sea-surface temperature.

## References

Allouche O, Tsoar A, Kadmon R 2006. Assessing the accuracy of species distribution models: prevalence, kappa and the true skill statistic (TSS). Journal of Applied Ecology 43:1223–1232 DOI 10.1111/j.1365-2664.2006.01214.x.

Bartoń K 2015. MuMIn: Multi-Model Inference. R package version 1.15.6.

Boria RA, Olson LE, Goodman SM, Anderson RP 2014. Spatial filtering to reduce sampling bias can improve the performance of ecological niche models. Ecological Modelling 275:73–77 DOI 10.1016/j.ecolmodel.2013.12.012.

Breiman L 2001. Random forests. Machine Learning 45:5–32.

Brown BE 1997. Coral bleaching: causes and consequences. Coral Reefs 16:S129–S138 DOI 10.1007/s003380050249.

Brown B, Dunne R, Goodson M, Douglas A 2002. Experience shapes the susceptibility of a reef coral to bleaching. Coral Reefs 21:119–126 DOI 10.1007/s00338-002-0215-z.

Cacciapaglia C, van Woesik R 2016. Climate-change refugia: shading reef corals by turbidity. Global Change Biology 22:1145–1154 DOI 10.1111/gcb.13166.

Donner SD 2009. Coping with commitment: projected thermal stress on coral reefs under different future scenarios. PLoS ONE 4:e5712 DOI 10.1371/journal.pone.0005712.

Donner SD 2011. An evaluation of the effect of recent temperature variability on the prediction of coral bleaching events. Ecological Applications 21:1718–1730 DOI 10.1890/10-0107.1.

Donner SD, Skirving WJ, Little CM, Oppenheimer M, Hoegh-Guldberg O 2005. Global assessment of coral bleaching and required rates of adaptation under climate change. Global Change Biology 11:2251–2265 DOI 10.1111/j.1365-2486.2005.01073.x.

Donner SD, Rickbeil GJM, Heron SF 2017. A new, high-resolution global mass coral bleaching database. PLoS ONE 12: e0175490 DOI 10.1371/journal.pone.0175490.

Dormann CF, Bierman SM, McPherson JM, Araujo MB, Bivand R, Bolliger J, Carl G, Davies RG, Hirzel A, Jetz W, Kissling WD, Kuhm I, Ohlemuller R, Peres-Neto PR, Reineking B, Schroder B, Schurr FM, Wilson R 2007. Methods to account for spatial autocorrelation in the analysis of species distributional data: a review. Ecography 30:609–628 DOI 10.1111/j.2007.0906-7590.05171.x.

Fabricius KE 2005. Effects of terrestrial runoff on the ecology of corals and coral reefs: review and synthesis. Marine Pollution Bulletin 50:125–146 DOI 10.1016/j.marpolbul.2004.11.028.

Geospatial Information Authority of Japan. 2015. Maps & Geospatial Information. Available at http://www.gsi.go.jp/ENGLISH/index.html (accessed at 25 May 2015).

Guest JR, Baird AH, Maynard JA, Muttaqin E, Edwards AJ, Campbell SJ, Yewdall K, Affendi YA, Chou LM 2012. Contrasting patterns of coral bleaching susceptibility in 2010 suggest an adaptive response to thermal Stress. PLoS ONE 7:e33353 DOI 10.1371/journal.pone.0033353.

Guinotte, J. M., Buddemeier, R. W., and Kleypas, J. A. 2003. Future coral reef habitat marginality: temporal and spatial effects of climate change in the Pacific basin. Coral Reefs 22:551–558 DOI 10.1007/s00338-003-0331-4.

Harii S, Hongo C, Ishihara M, Ide Y, Kayanne H 2014. Impacts of multiple disturbances on coral communities at Ishigaki Island, Okinawa, Japan, during a 15 year survey. Marine Ecology Progress Series 509:171–180 DOI 10.3354/meps10890.

Heron SF, Liu G, Rauenzahn JL, Christensen TRL, Skirving WJ, Burgess TFR, Eakin CM, Morgan JA 2014. Improvements to and continuity of operational global thermal stress monitoring for coral bleaching. Journal of Operational Oceanography 7:3–11 DOI 10.1080/1755876X.2014.11020154.

Hijmans RJ, Phillips SJ, Leathwick J, Elith J 2017. dismo: Species Distribution Modeling. version 1.1-4. Available at https://cran.r-project.org/web/packages/dismo/index.html. (accessed 26 Sep. 2017)

Hoegh-Guldberg O 1999. Climate change, coral bleaching and the future of the world’s coral reefs. Marine and Freshwater Research 50:839–866 DOI 10.1071/MF99078.

Hongo C, Yamano H 2013. Species-specific responses of corals to bleaching events on anthropogenically turbid reefs on Okinawa Island, Japan, over a 15-year period (1995–2009). PLoS ONE 8: e60952 DOI 10.1371/journal.pone.0060952.

Jones G, Curran M, Swan H, Deschaseaux E 2017. Dimethylsulfide and coral bleaching: links to solar radiation, low level cloud and the regulation of seawater temperatures and climate in the Great Barrier Reef. American Journal of Climate Change 6:328–359 DOI 10.4236/ajcc.2017.62017.

JPL MUR MEaSUREs Project. 2015. GHRSST Level 4 MUR Global Foundation Sea Surface Temperature Analysis (v4.1). Ver. 4.1. PO.DAAC, CA,USA. Available at http://dx.doi.org/10.5067/GHGMR-4FJ04. (accessed 21 September 2016)

Kayanne H 2017. Validation of degree heating weeks as a coral bleaching index in the northwestern Pacific. Coral Reefs 36:63–70 DOI 10.1007/s00338-016-1524-y.

Kayanne H, Suzuki R, Liu G 2017. Bleaching in the Ryukyu Islands in 2016 and associated Degree Heating Week threshold. Galaxea, Journal of Coral Reef Studies 19:17–18 DOI 10.3755/galaxea.19.1_17.

Kleypas JA, McManus JW, Meñez LAB 1999. Environmental limits to coral reef development: where do we draw the line? American Zoologist 39:146–159 DOI 10.1093/icb/39.1.146.

Kleypas JA, Danabasoglu G, Lough JM 2008. Potential role of the ocean thermostat in determining regional differences in coral reef bleaching events. Geophysical Research Letters 35:L03613 DOI 10.1029/2007GL032257.

Liaw A, Wiener M 2002. Classification and Regression by randomForest. R News 2:18–22.

Liu G, Strong AE, Skirving WJ 2003. Remote sensing of sea surface temperatures during 2002 barrier reef coral bleaching. EOS, Transactions of the American Geophysical Union 84:137–141 DOI 10.1029/2003EO150001.

Liu C, Berry PM, Dawson TP, Pearson RG 2005. Selecting thresholds of occurrence in the prediction of species distributions. Ecography 28:385–393 DOI 10.1111/j.0906-7590.2005.03957.x.

Liu G, Heron SF, Eakin CM, Muller-Karger FE, Vega-Rodriguez M, Guild LS, De La Cour JL, Geiger EF, Skirving WJ, Burgess TFR, Strong AE, Harris A, Maturi E, Ignatov A, Sapper J, Li J, Lynds S 2014. Reef-scale thermal stress monitoring of coral ecosystems: New 5-km global products from NOAA Coral Reef Watch. Remote Sensing 6:11579–11606 DOI 10.3390/rs61111579.

Liu G, Skirving WJ, Geiger EF, De La Cour JL, Marsh BL, Heron SF, Tirak KV, Strong AE, Eakin CM 2017. NOAA Coral Reef Watch's 5km Satellite Coral Bleaching Heat Stress Monitoring Product Suite Version 3 and Four-Month Outlook Version 4. Reef Encounter 32: 39–45. Available at http://coralreefs.org/wp-content/uploads/2014/03/Reef-Encounter-August-2017-FINAL-Lo-Res.pdf (accessed 2 October 2017)

Maina J, Venus V, McClanahan TR, Ateweberhan M 2008. Modelling susceptibility of coral reefs to environmental stress using remote sensing data and GIS models. Ecological Modeling 212:180–199 DOI 10.1016/j.ecolmodel.2007.10.033.

Maina J, McClanahan TR, Venus V, Ateweberhan M, Madin J 2011. Global Gradients of Coral Exposure to Environmental Stresses and Implications for Local Management. PLoS ONE 6:e23064 DOI 10.1371/journal.pone.0023064.

Manel S, Williams HC, Ormerod SJ 2001. Evaluating presence-absence models in ecology: the need to account for prevalence. Journal of Applied Ecology 38:921–931 DOI 10.1046/j.1365-2664.2001.00647.x.

Maynard JA, Anthony KRN, Marshall PA, Masiri I 2008. Major bleaching events can lead to increased thermal tolerance in corals. Marine Biology 155:173–182 DOI 10.1007/s00227-008-1015-y.

McWilliams JP, Cote IM, Gill JA, Sutherland WJ, Watkinson AR. 2005. Accelerating impacts of temperature-induced coral bleaching in the Caribbean. Ecology 86:2055–2060 DOI 10.1890/04-1657.

McClanahan TR 2014. Decadal coral community reassembly on an African fringing reef. Coral Reefs 33:939–950 DOI 10.1007/s00338-014-1178-6.

McClanahan TR, Maina F, Ateweberhan M 2015. Regional coral responses to climate disturbances and warming is predicted by multivariate stress model and not temperature threshold metrics. Climatic Change 131:607–620 DOI 10.1007/s10584-015-1399-x.

Nakamura T, van Woesik R. 2001. Water-flow rates and passive diffusion partially explain differential survival of corals during the 1998 bleaching event. Marine Ecology Progress Series 212:301–304 DOI 10.3354/meps212301.

Namizaki N, Yamano H, Suzuki R, Oohori K, Onaga H, Kishimoto T, Sagawa T, Machida Y, Yasumura S, Satoh T, Shigiya T, Shibata T, Tsuchikawa M, Miyamoto Y, Harukawa K, Hirate K, Furuse K, Hokoyama K, Yamakawa Y, Wagatsuma T 2013. The potential of citizen monitoring programs for marine areas: activities of the two-year Sango (Coral) Map Project. Galaxea, Journal of Coral Reef Studies 15:391–395 DOI 10.3755/galaxea.15.391.

Okinawa Prefecture 2017. Development of Methods for Coral Seedling Production by Sexual Reproduction. Department of Environment, Okinawa Prefectural Government.

Oliver JK, Berkelmans R, Eakin CM 2009. Coral Bleaching in space and time. In: van Oppen MJH, Lough JM, ed. Coral Bleaching: Patterns, Processes, Causes and Consequences: Springer-Verlag Berlin, 21–39.

Oxenford HA, Vallés H 2016. Transient turbid water mass reduces temperature-induced coral bleaching and mortality in Barbados. PeerJ 4:e2118 DOI 10.7717/peerj.2118.

Palumbi SR, Barshis DJ, Traylor-Knowles N, Bay RA 2014. Mechanisms of reef coral resistance to future climate change. Science 344:895–898 DOI 10.1126/science.1251336.

R Core Team 2017. R: A language and environment for statistical computing. R Foundation for Statistical Computing, Vienna, Austria; https://www.R-project.org/ (accessed 30 June 2017).

Strong AE, Liu G, Kimura T, Yamano H, Tsuchiya M, Kakuma S, van Woesik R 2002. Detecting and monitoring 2001 coral reef bleaching events in Ryukyu Islands, Japan using satellite bleaching HotSpot remote sensing technique. Geoscience and Remote Sensing Symposium 2002, IEEE International 1:237–239 DOI 10.1109/IGARSS.2002.1024998.

Staub F, Chhay L 2009. International Year of the Reef 2008 The Year in Review; https://docs.lib.noaa.gov/noaa_documents/CoRIS/IYOR/IYOR_2008.pdf (accessed 17 October 2017)

Tabor K, Williams JW 2010. Globally downscaled climate projections for assessing the conservation impacts of climate change. Ecological Applications 20:554–565 DOI 10.1890/09-0173.1.

Torda G, Donelson JM, Aranda M, Barshis DJ, Bay L, Berumen ML, Bourne DG, Cantin N, Foret S, Matz M, Miller DJ, Moya A, Putnam HM, Ravasi T, van Oppen MJH, Thurber RV, Vidal-Dupiol J, Voolstra CR, Watson S-A, Whitelaw E, Willis BL, Munday PL 2017. Rapid adaptive responses to climate change in corals. Nature Climate Change 7:627–636 DOI 10.1038/nclimate3374.

Tupper M, Tan M, Tan S, Radius M, Abdullah S 2011. ReefBase: a global information system on coral reefs; http://www.reefbase.org (accessed 15 August 2017).

van Hooidonk R, Huber M 2009. Quantifying the quality of coral bleaching predictions. Coral Reefs 28:579–587 DOI 10.1007/s00338-009-0502-z.

Welle PD, Small MJ, Doney SC, Azevedo IL 2017. Estimating the effect of multiple environmental stressors on coral bleaching and mortality. PLoS ONE 12:e0175018 DOI 10.1371/journal.pone.0175018.

West JM, Salm RV 2003. Resistance and resilience to coral bleaching: Implications for coral reef conservation and management. Conservation Biology 17:956–967 DOI 10.1046/j.1523-1739.2003.02055.x.

Wooldridge SA, Done TJ 2009. Improved water quality can ameliorate effects of climate change on corals. Ecological Applications 19:1492–1499 DOI 10.1890/08-0963.1.

Wu L, Cai W, Zhang L, Nakamura H, Timmermann A, Joyce T, McPhaden MJ, Alexander M, Qiu B, Visbeck M, Chang P, Giese B 2012. Enhanced warming over the global subtropical western boundary currents. Nature Climate Change 2:161–166 DOI 10.1038/nclimate1353.

Yara Y, Oshima K, Fujii M, Yamano H, Yamanaka Y, Okada N 2011. Projection and uncertainty of the poleward range expansion of coral habitats in response to sea surface temperature warming: a multiple climate model study. Galaxea, Journal of Coral Reef Studies 13:11–20 DOI 10.3755/galaxea.13.11.

Yee SH, Santavy DL, Barron MG 2008. Comparing environmental influences on coral bleaching across and within species using clustered binomial regression. Ecological Modelling 218:162–174 DOI 10.1016/j.ecolmodel.2008.06.037.

